# Overcoming steric inhibition of antibody-dependent phagocytosis with tall adhesions

**DOI:** 10.64898/2026.01.09.698749

**Authors:** Aaron M. Joffe, Aymeric Chorlay, Jared Huzar, Jaffar Hasnain, Phillip L. Geissler, Daniel A. Fletcher

## Abstract

Macrophages recognize and phagocytose opsonized target cells, including those coated with IgG antibodies. This process relies on binding of IgG to Fcγ receptors (FcγR) expressed on the macrophage surface, resulting in formation of a phagocytic synapse. Since the surface of both macrophages and target cells are densely packed with macromolecules of diverse sizes, most of which are not directly involved in phagocytic signaling, it is possible for tall ‘bystander’ proteins to sterically interfere with FcγR engagement. Here, we use cell-like target particles to show that bystander proteins can inhibit phagocytosis by blocking synapse formation. We then demonstrate that adding a tall binding protein to the target particle can overcome inhibition by the crowded environment and substantially recover phagocytosis, a process we call kinetic enhancement. Using a cell-free system of giant unilamellar vesicles and synthetic binders, we demonstrate that kinetic enhancement is a tunable feature of interface formation that can determine whether short binders engage, and we present theory and computer simulations to explain the nonmonotonic dependence of phagocytosis on tall binding protein surface density. These findings point to a strategy for overcoming surface crowding on phagocytic targets by re-engineering transition states with tall adhesion proteins, one that could be used to promote short receptor binding at other cell-cell junctions.

**Significance Statement:** Macrophages contribute to our immune defenses by phagocytosing pathogens and diseased cells. To accomplish this, they must first establish close contacts between receptors on their membranes and antibodies or other ligands decorating target cells. However, macrophage binding to the target can be disrupted by the presence of tall neighboring proteins and glycans—‘bystander’ molecules—that sterically prevent the two surfaces from coming into close contact. Counterintuitively, this inhibition can be overcome by the addition of even taller binding proteins between the macrophage and target cell, albeit at low concentrations. Using live cell and in vitro experiments, theory, and computer simulations, we show that tall binders can promote close contact that enables phagocytosis, even in the presence of bystander proteins that would normally block close contact.

## Introduction

Cell surfaces are crowded environments consisting of thousands of membrane-bound molecules per square micron that extend beyond the plasma membrane by tens to hundreds of nanometers, or more [1]. At sufficient densities and glycosylation levels, these molecules create a steric barrier that can inhibit close contact and subsequent receptor binding between cells, especially for short binders. Many cells, like macrophages, rely on close contact to carry out their function. During an immune response, macrophages are critical actors that recognize and clear pathogens and foreign objects by phagocytosis, while also initiating broad-scale immune responses by secreting inflammatory cytokines and presenting antigens to adaptive immune cells [2–4]. Although macrophages recognize many classical pathogens using a diverse set of pattern recognition receptors on their surface, detection of novel targets, including highly mutagenic pathogens and host-derived cancer cells, often involves recognition of antibodies through Fc receptors. In one version of this detection strategy, IgG antibodies serve as adaptor molecules that bind to unique antigen epitopes on the pathogen surface via their Fab regions and make the target susceptible to recognition by Fcγ receptors (FcγRs) FcγRs through binding to the Fc region of IgG. This mechanism of phagocytosis, known as antibody dependent cellular phagocytosis (ADCP), has been harnessed to target and clear cancer cells during treatment with antibody-based immunotherapies [5,6].

Therapeutic strategies that maximize activation and minimize inhibition of macrophages are being pursued through multiple avenues. The most direct routes are developing specific targeting antibodies that enhance activation signals or high-affinity blocking antibodies that disrupt inhibitory checkpoints [7]. In previous work, we studied the effect of antigen height on ADCP and found that antibodies targeting antigens with short extracellular domains (< 10 nm) were more efficient at triggering phagocytosis due to the requirement for size-dependent segregation of CD45 phosphatase molecules from the phagocytic interface [8]. We also showed that adding CD47 to the IgG-coated surface inhibited phagocytosis in a ratiometric way [9]. In these studies, we used reconstituted cell-like target particles with only the proteins of interest—activating IgG-bound antigens and/or inhibitory CD47—that allowed us to isolate the role of specific molecular players in phagocytosis. However, it remains unclear how the complex surface of a real target cell, which contains many molecules not directly involved in phagocytic signaling, might affect synapse formation and FcγR engagement.

Short antigens typically targeted by therapeutic antibodies on tumor cells constitute only a small subset of all surface proteins and are surrounded by a dense array of tall non-antigenic proteins [10,11], including tall, heavily glycosylated proteins such as MUC1, P-selectin, and CEACAM5. One might expect that the presence of this forest of non-antigenic proteins, which we refer to as ‘bystander’ proteins, could act as a steric barrier to IgG-FcγR binding [12,13]. This idea is consistent with previous simulations [14] and the use of poly(ethyleneglycol) (PEG) polymer brushes to block molecular adsorption to surfaces [15,16]. Steric barriers to phagocytosis would be most extreme in the case of IgG bound to short antigens on the surface of cancer cells, especially given that cancer cells have been shown to upregulate the expression of tall glycoproteins [17,18]. Indeed, cell surface crowding has been demonstrated to impose an energy penalty on the scale of 1-3k_B_T for an IgG in solution binding to buried receptors [12]. In contrast to better studied inhibitory checkpoints such as CD47-SIRPα [9], steric barriers created by bystander proteins represent a potentially potent and ubiquitous inhibitory mechanism that is poorly understood and for which therapeutic strategies to overcome it are lacking.

In this work, we use cell-like reconstituted target particles to explore the effect of bystander protein size and density on ADCP. Using live cell imaging, we quantify bystander-mediated inhibition of phagocytosis as a function of bystander protein height and abundance. Counterintuitively, we find that the addition of a tall macrophage binding protein (P-selectin) to target surfaces can restore efficient phagocytosis. To test if this phenomenon, which we call ‘kinetic enhancement’, requires active cell processes or is a property of membrane interface formation in the presence of blocking proteins, we reconstitute interface formation using giant unilamellar vesicles, DNA binders, and purified bystander proteins. We observe similar effects of bystanders and long binders as in our macrophage experiments, suggesting that steric inhibition of interface formation by bystander proteins and subsequent recovery with long binders is a general phenomenon governed by equilibrium thermodynamics and potentially applicable to other cell-cell interfaces.

To clarify the microscopic mechanisms underlying kinetic enhancement, we use theory and computer simulations to show that bystander-mediated thermodynamic barriers to phagocytosis can be crossed in a reasonable time only if the corresponding transition state involves strong local deformation in membrane shape—a dimple of IgG-FcγR binding that can progressively offset the entropic cost of excluding bystanders from the nascent synapse. Tall adhesion proteins stabilize a distinct but similarly dimpled state, with the potential to greatly increase the rate of engagement. Their binding, however, also establishes an undimpled intermediate state that becomes deeply metastable at high density (or strong binding) of the tall adhesion proteins, with the potential to greatly impede close contact.

These findings add to previous understanding of phagocytic synapse formation and FcγR engagement, showing that while targeting IgG antibodies to short antigens may most efficiently trigger signaling once the phagocytic synapse is formed, tall non-binding proteins around those short antigens can sterically impede formation of the phagocytic synapse in the first place. We demonstrate that the transition states introduced by bystander proteins can be engineered with additional tall binders to facilitate barrier crossing, an approach that cells may have evolved to promote engagement of short ligand-receptor pairs in physiological contexts and one that could be co-opted for therapeutic purposes.

## Results

### Non-binding bystander proteins can inhibit antibody-dependent phagocytosis

In order to determine if bystander proteins can inhibit macrophage antibody-dependent phagocytosis, we created reconstituted target particles to model antibody opsonized cells and added bystander proteins with controlled densities and heights using a previously described method [19]. Briefly, each particle consists of a glass bead coated with a fluid lipid bilayer to model the fluid plasma membrane of a target cell, with His-tagged proteins of interest directly coupled to Ni-NTA lipids in the bilayer and monoclonal IgG antibodies bound to specific antigens (Figure 1A). In our experiments, biotinylated lipids were incorporated into the lipid bilayer so that anti-biotin IgG could be bound directly to the particle surface. This configuration models an antibody bound to a short antigen, which we previously found to efficiently trigger phagocytic signaling due to size-based segregation of the CD45 phosphatase from the phagocytic interface [8]. All experimental conditions used a fixed surface density of anti-biotin IgG of 200 per µm^-2^ as described in [8].

**Figure 1:**
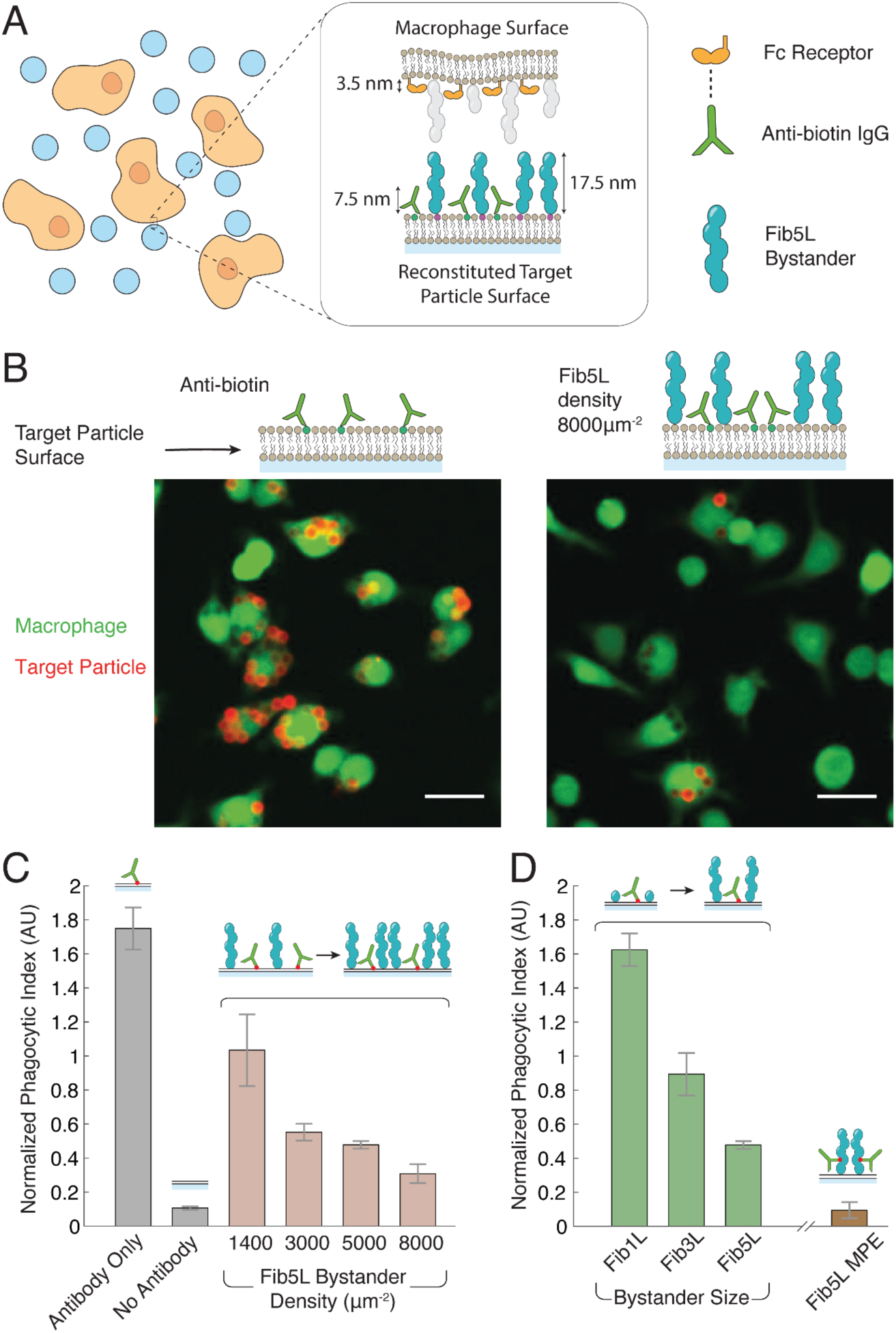
Bystander proteins on target membranes can inhibit antibody-mediated cellular phagocytosis. **(A)** Macrophages are incubated with reconstituted phagocytic targets to assay phagocytosis. Important proteins involved in phagocytic interface formation are illustrated along with their respective sizes. **(B)** Fluorescence images showing macrophages incubated with reconstituted targets containing anti-biotin IgG with or without the addition of Fib5L bystander. Scale bars: 20 µm. **(C)** Reconstituted targets displaying both anti-biotin IgG and Fib5L bystander protein were incubated with macrophages. The first two bars represent a positive control (targets displaying only anti-biotin IgG) and a negative control (targets displaying no anti-biotin IgG) respectively. Error bars represent standard error for >3 replicates of the same condition. **(D)** Reconstituted targets displaying both anti-biotin IgG and Fib5L, Fib3L, or Fib1L bystander protein were incubated with macrophages. The first two bars represent a positive control (targets displaying only anti-biotin IgG) and a negative control (targets displaying no anti-biotin IgG) respectively. Error bars represent standard error for >3 replicates of the same condition.

To test the effect of bystander protein size and surface density on ADCP, we used synthetic proteins made of repeated fibronectin-like domains [20] (Fibcons) that were conjugated to Ni-NTA lipids in the bilayer. The Fibcon proteins used in these experiments contained 1, 3, or 5 domain repeats (Fib1L, Fib3L and Fib5L, respectively), corresponding to fully extended protein heights of 3.5 nm, 10.5 nm, and 17.5 nm. The density of bystander proteins displayed on the surface of target particles was controlled by changing the amount of Ni-NTA lipids in the bilayer. Absolute densities of Fibcon proteins were measured with flow cytometry using fluorescently-labeled versions of the proteins calibrated to beads with known fluorophore numbers (Figure S1).

As a first test of bystander-mediated inhibition, we compared phagocytosis of target particles displaying only anti-biotin antibodies with target particles displaying both anti-biotin antibodies and bystander proteins. We began with Fib5L as the bystander because its height of 17.5 nm is closest to the average cell surface protein height of 15 nm [21]. The Fib5L bystander proteins were displayed at a density of 8,000 µm^-2^, which is below reported bulk cellular surface densities of approximately 20,000 µm^-2^ [22]. The target particles were incubated in a 96-well plate with RAW 264.7 macrophage-like cells for 30 min at 37 °C, followed by washing of excess particles from the well. Fluorescent imaging of the macrophages after incubation revealed dramatically less phagocytosis for conditions in which bystander proteins were present compared to those without bystander proteins (Figure 1B).

We next quantified phagocytosis for a range of Fib5L densities and heights, alongside a negative control displaying no anti-biotin IgG and a positive control displaying only anti-biotin IgG with no bystander proteins. Defining the average value of internalized target fluorescence per macrophage as the ‘phagocytic index’, we observed a steady decrease in phagocytosis with increasing density of Fib5L bystanders (Figure 1C). To determine the effect of bystander protein height on phagocytosis, we then fixed the density of bystander proteins to be 5,000 µm^-2^ but varied the height by using Fib1L, Fib3L, or Fib5L. Quantification of phagocytic indices for these conditions showed that phagocytosis decreased as bystander height increased (Figure 1D).

Since short antigens are more effective at driving phagocytosis but long bystanders can block phagocytosis, we wondered whether the tall antigens could be overcome by targeting antibodies to membrane-proximal epitopes on the tall antigens. To test this, we converted Fib5L into an antigen by biotinylating the membrane-proximal C-terminus of Fib5L, added anti-biotin IgG to Fib5L at a surface density of 8,000 µm^-2^, and quantified phagocytosis. Even though antibody density is significantly higher than for the case of biotinylated lipids, we find that phagocytosis is still blocked (Figure 1D). This data suggests that targeting the membrane-proximal epitope of a tall antigen will not enhance phagocytosis since the tall antigen effectively blocks FcγR engagement, similar to a bystander protein at high surface densities.

When antibodies are attached to a membrane-distal epitope of antigens with increasing height (biotinylated Fib1L, Fib3L, and Fib5L) and also surrounded by tall bystander proteins (Fib5L), we find that phagocytosis modestly recovers only for biotinylated Fib3L, the tallest antigen that is still capable of creating a membrane gap, when bound to FcγR, that excludes CD45 (Figure S2). Taken together, these results indicate that bystander proteins can sterically inhibit phagocytosis of antibody-opsonized particles.

### Bystander proteins impede FcγR engagement and dynamics at the phagocytic interface

To confirm that bystander proteins are inhibiting phagocytosis by interfering with phagocytic interface formation and IgG-FcγR binding, we directly imaged interface formation using total internal reflection fluorescence (TIRF) microscopy. To do this, we created supported lipid bilayers (SLBs) on glass coverslips using the same IgG and bystander protein compositions used to test phagocytosis with glass beads. We then dropped macrophages onto the SLBs coated with IgG and bystander proteins and tracked fluorescent anti-biotin IgG dynamics at the interface after engagement with macrophage FcγRs (Figure 2A). We also captured images using Reflection Interference Contrast Microscopy (RICM) to quantify the contact area of each macrophage, regardless of its ability to bind and cluster antibody.

**Figure 2:**
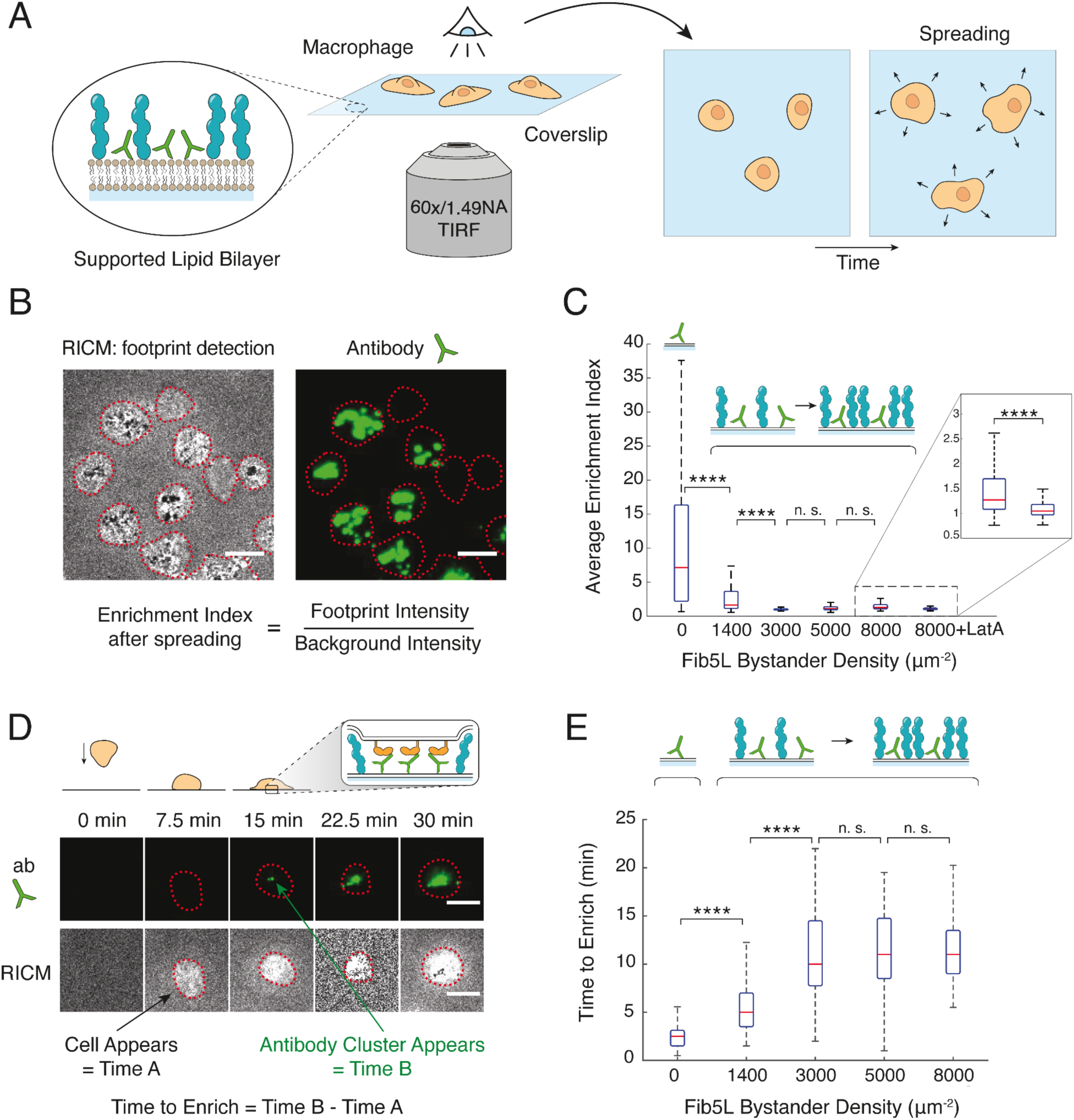
Bystander proteins sterically block antibody engagement with macrophage Fc receptors. **(A)** Schematic view of TIRF live-cell imaging experiments to visualize the phagocytic interface. **(B)** TIRF images of fluorescently labeled anti-biotin IgG were captured alongside RICM images of macrophages. The enrichment index was quantified by measuring the fluorescence intensity within each macrophage interface (masked in RICM channel) and dividing by the background fluorescence intensity. Images were captured after a 30-minute incubation to represent the equilibrium state. Scale bars: 10 µm **(C)** Enrichment indices for macrophages interacting with SLBs displaying different densities of Fib5L bystander protein. Inset shows a comparison between macrophages with and without the addition of LatA for a Fib5L bystander density of 8,000 µm^-2^. Each condition represents values collected from > 35 macrophages per day, across 3 days (all data combined). **** represents a p value < 0.00001 resulting from a two-sample Student’s t-test. **(D)** TIRF images of fluorescently labeled anti-biotin IgG were captured alongside RICM images of macrophages every 30 seconds for a total of 30 minutes. The time to enrich was measured by subtracting the time at which a macrophage first appeared in the RICM channel from the time at which anti-biotin IgG clustering was first observed in the TIRF channel. Scale bars: 10 µm. **(E)** The time to enrich for macrophages interacting with SLBs displaying different densities of Fib5L bystander protein. Each condition represents values collected from > 20 macrophages per day, across 3 days (all data combined). **** represents a p value < 0.00001 resulting from a two-sample Student’s t-test.

First, we characterized the amount of anti-biotin IgG bound to FcγR at the interface between each macrophage and the SLB after 30 minutes of incubation, which is the point at which interfaces reached a quasi-steady state. To do this, we used RICM to identify each macrophage in the field of view and created a mask around its area. Next, we quantified an ‘enrichment index’ (EI) by measuring the average IgG fluorescence intensity within each macrophage mask and dividing it by the average IgG intensity outside of the mask (Figure 2B). EI acts as a metric for interface formation by quantifying the amount of antibody bound at the phagocytic interface for each macrophage.

We observed a substantial decrease in EI for cells as the density of bystander Fib5L was increased, dropping from a median value of 7.1 down to 1.6 in the presence of bystander proteins at a surface density of 1,400 µm^-2^ (Figure 2C). This observation is consistent with bystander proteins inhibiting phagocytosis by blocking phagocytic interface formation. Interestingly, macrophages treated with Latrunculin A, which depolymerizes the actin cytoskeleton, showed an even further reduction in interface formation in the presence of dense Fib5L bystanders when compared to untreated macrophages (Figure 2C, inset). This suggests that the actin cytoskeleton is playing an important role overcoming the steric barrier to interface formation presented by bystander proteins, likely through active probing of surroundings with protrusive structures [23,24].

To further characterize the effect of bystander proteins on IgG-FcγR binding, we next used time-lapse imaging to observe the dynamics of interface formation. Using the same conditions as in the enrichment index measurements, we captured images of macrophages interacting with the SLB every 30 seconds for 30 minutes. For each macrophage that formed antibody clusters within that timeframe, we measured the time to enrich (*t_enrich_*) as the difference between the time at which the first IgG cluster appeared in TIRF and the time at which the macrophage first settled on the surface in RICM (Figure 2D). The median value for *t_enrich_* was 2.5 minutes for macrophages binding to SLBs displaying only IgG. However, when bystander Fib5L was added at a surface density of 1,400 ±200 µm^-2^, this value doubled to become 5 minutes. For bystander Fib5L surface densities of 3,000 ±500 µm^-2^ and above, *t_enrich_* increased to 10 minutes (Figure 2E).

This observation indicates that bystander proteins not only lessen the equilibrium number of IgG-FcγR complexes formed at the phagocytic interface at long times but also decrease the rate at which clusters form. Considering that average speeds of tissue resident macrophages have been measured to be on the order of 1 µm/minute [25], the average contact time between a macrophage and a potential target cell would last less than approximately 10 minutes (assuming a stationary target cell with a diameter of 10 µm). The measured shift in *t_enrich_* from 2.5 minutes to 10 minutes due to the increased surface density of bystander proteins therefore has the potential to significantly alter the likelihood of phagocytic synapse formation and signaling over physiologically relevant timescales.

### P-selectin, a tall binding protein, can partially rescue phagocytosis by overcoming bystander-mediated blocking

Our results indicate that the presence of bystander proteins can substantially inhibit the engagement of short binding partners at the macrophage-target interface. In addition to these non-adhesive bystander proteins, natural cell-cell interfaces are also populated by a variety of long adhesion proteins. Like bystanders, these species must be excluded from an interface to permit close contact and IgG-FcγR binding, carrying an even steeper thermodynamic cost due to the disruption of favorable contacts or the exclusion of bound pairs of binders. Unlike bystanders, however, tall binders stabilize cell-cell attachments from which a phagocytic synapse could form with relatively modest organizational changes at the membrane interface, namely membrane deformation leading to short binder engagement and exclusion of both the tall binder and bystander proteins. We hypothesized that such tall binders could, under some circumstances, alleviate bystanders’ kinetic inhibition of phagocytic synapse formation and restore phagocytosis.

We experimentally tested this idea by adding P-selectin to target particles, in addition to Fib5L bystander proteins and IgG bound to biotinylated lipids. P-selectin is typically expressed by activated platelets and endothelial cells and is known to bind to its ligand PSGL-1 on leukocytes and leukocyte-derived cells, including macrophages [26]. It is a rigid protein with a measured height of 38 nm [27,28] that binds to PSGL-1 at its distal end. We chose P-selectin because (i) it is longer than the Fibcon-based bystander proteins, enabling easy binding, (ii) it binds to the macrophage without directly activating or inhibiting phagocytic signaling, and (iii) its affinity for PSGL-1 is low [29] so that binding of the FcγR to IgG [30] is still the favored state at equilibrium. Interestingly, P-selectin is known to be critical for leukocyte transmigration, which involves leukocyte rolling along the endothelial lining, arrest, and ultimately the formation of stable integrin binding events that initiate transmigratory signaling [31,32]. We wondered if P-selectin present on the surface of a phagocytic target could—mirroring its role in transmigration—yield kinetic enhancement of IgG-FcγR binding.

We first investigated whether P-selectin displayed on a target particle’s surface would lead to increased binding between macrophages and targets. We created target particles with IgG antibody, Fib5L bystander protein, and purified P-selectin containing a C-terminal His-tag that was conjugated to Ni-NTA lipids in the target particle bilayer (Figure 3A). As in all previous experiments, the anti-biotin IgG density was fixed at a surface density of 200 µm^-2^. The total combined surface density of Fib5L bystander plus P-selectin was set at 8,000 µm^-2^ by fixing the total concentration of proteins added to the target particles, while the relative molar percent of P-selectin was varied from 0% to 100%.

**Figure 3:**
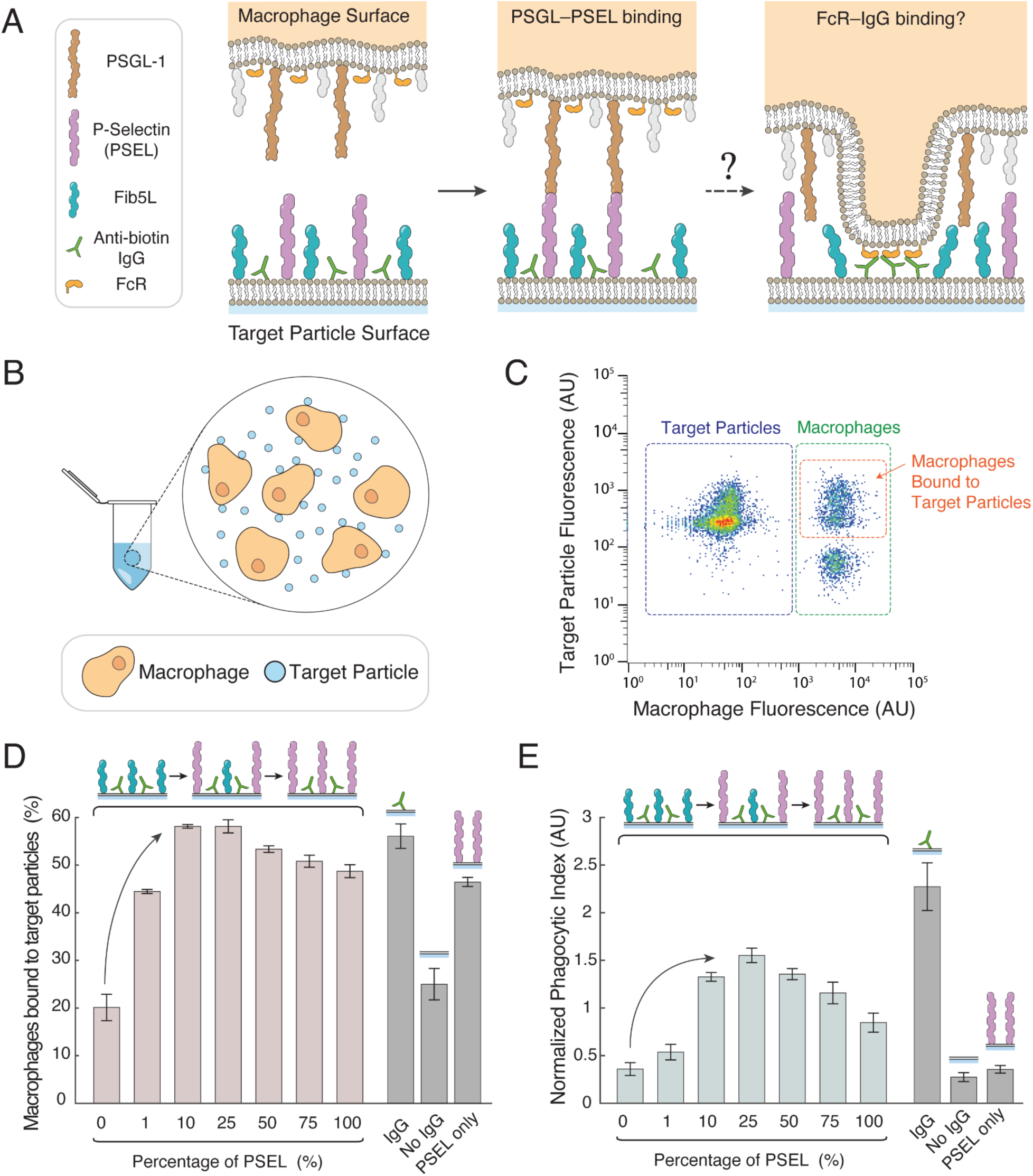
Addition of tall binding proteins to the target membrane can rescue phagocytosis of bystander-blocked particles. **(A)** Illustration of a reconstituted target particle surface containing anti-biotin IgG, Fib5L bystander proteins, and tall binding protein P-selectin. Upon binding of P-selectin to its ligand PSGL-1 on the macrophage surface, the interface can remain in the metastable state with P-selectin bound to PSGL-1 or form a phagocytic interface by engagement of IgG with the macrophage FcγR, leading to phagocytosis. **(B)** To assay interface formation, macrophages are incubated with reconstituted target particles at 4°C, which inhibits phagocytosis. **(C)** Flow cytometry shows distinct populations of target particles and macrophages. Interface formation is quantified by measuring the percentage of macrophages that contain target particle fluorescence. **(D)** Quantification of interface formation for target particles displaying varying amounts of anti-biotin IgG, Fib5L bystander protein, and P-Selectin (PSEL). The first two bars represent a positive and negative control, respectively, and give the dynamic range of the measurement. The addition of P-Selectin on target particles leads to an increase in interface formation above levels containing only Fib5L bystander (indicated by arrow). **(E)** Quantification of phagocytosis of target particles displaying the same surface molecules as in D. The addition of P-Selectin on the target surface leads to an increase in phagocytosis above levels containing only Fib5L bystander (indicated by arrow).

To determine if interface formation occurs in the absence of any phagocytosis, we first incubated fluorescently labeled macrophages and targets in tubes under constant rotation at 4 °C, a temperature at which macrophages can bind phagocytic targets but are unable to phagocytose them (Figure 3B). We ran the contents of the tube through a flow cytometer and plotted fluorescence intensities from the target particles and macrophages on orthogonal axes (Figure 3C). Using this target binding assay, the macrophage population could be easily distinguished from the target particle population, and the macrophages showed two distinct sub-populations: those with target fluorescence and those with no target fluorescence. Because the targets could not be internalized by the macrophages, any macrophages showing target particle fluorescence must have had targets bound to their surface.

To compare target binding under different conditions, we calculated the percentage of macrophages that contained targets bound to their surface. We first measured binding of targets displaying only anti-biotin IgG on their surface and those without IgG, which represent our positive and negative controls and show the dynamic range of this measurement (Figure 3D). We next tested target particles displaying only P-selectin and confirmed that it does indeed lead to macrophage binding (Figure 3D). Finally, we measured binding to targets containing anti-biotin IgG, Fib5L bystander, and P-selectin over a range of concentrations. The results showed that the addition of P-selectin increased target binding to similar levels as the positive control condition. The maximum binding levels were reached with P-selectin at around 10% to 25%, which corresponds to a P-selectin surface density of 800 to 2,000 µm^-2^. Interestingly, these densities are only slightly higher than the surface densities of P-selectin on activated platelets of 450 to 750 µm^-2^, calculated from previous measurements of P-selectin surface expression [33–35] and mean platelet diameter [36].

After confirming that P-selectin increases adhesion between macrophages and target particles, we asked whether the presence of P-selectin also increases phagocytosis in the presence of bystander proteins by performing phagocytosis assays for each of the same conditions tested in the binding assay. Remarkably, we observed a significant increase in phagocytosis for conditions containing P-selectin alongside Fib5L bystander proteins (Figure 3E). Maximum phagocytosis occurred when P-selectin accounted for 25% of the bystander proteins on the surface, which is consistent with maximum macrophage-particle adhesion in our binding assay. We observed phagocytosis reaching 70% of positive control levels—a 4-fold increase in phagocytosis compared to phagocytosis of targets containing only anti-biotin IgG and Fib5L bystander proteins. Phagocytosis sharply drops off for P-selectin levels above 25%, presumably due to the added steric barrier of the long adhesions. Importantly, targets displaying only P-selectin on their surface lead to no phagocytosis (Figure 3E) even though they promote increased binding (Figure 3D). This confirms that P-selectin on its own has no pro-phagocytic signaling ability and that the increase in phagocytosis comes from kinetic enhancement of IgG-FcγR binding mediated by the long adhesions.

### Tall binders enable short binders to overcome bystander-mediated blocking in a minimal synthetic system

Our results indicate that a tall binder, P-selectin, can play a crucial role in restoring phagocytosis in the presence of bystander proteins. To test whether this phenomenon is the result of equilibrium membrane interface formation in the absence of active cellular processes, we reconstituted a minimal membrane system mimicking the interaction between macrophages and target particles. Using giant unilamellar vesicles (GUVs) functionalized with high-affinity short ligands (representing antibody-FcγR complexes), low-affinity tall ligands (mimicking P-selectin-PSGL1 complexes), and bystander proteins (Figure 4A), we investigated whether bystander proteins could be excluded and close membrane contacts formed by the right combination of short and tall binders.

**Figure 4:**
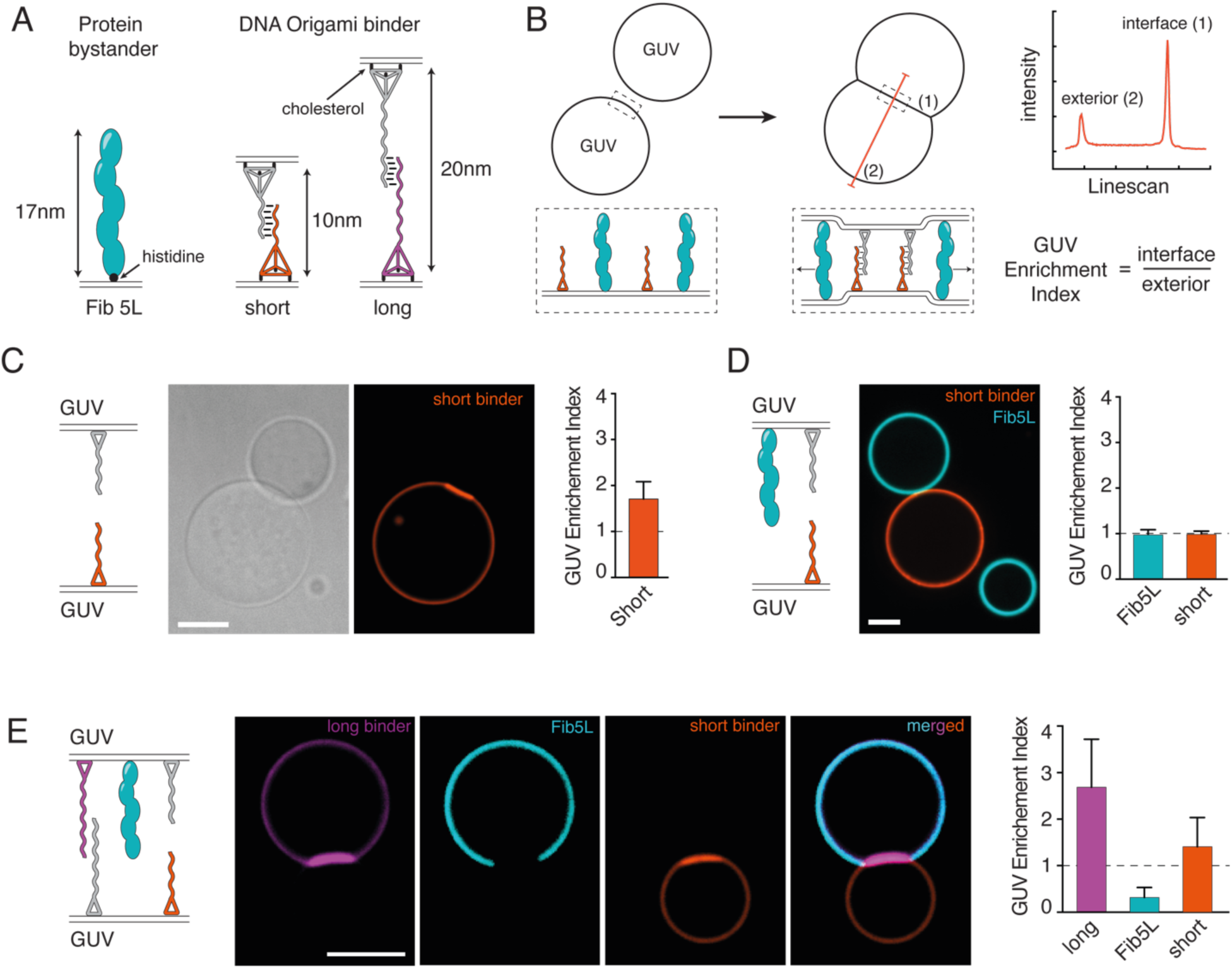
Tall binders help short binders overcome bystander barriers in reconstituted adhesion systems. **(A)** Purified bystander proteins (Fib 5L ∼17 nm) and complementary DNA origami binding pair (short binder pair ∼10 nm, and long binder pair ∼20 nm), are used to test if tall binders help short binders overcome bystander screening during membrane interface formation. **(B)** When GUVs containing DNA binders and bystander proteins interact, membrane interfaces form through DNA origami dimerization. Fib5L proteins attach to GUV membranes via His-tags on DGS-NTA(Ni), while DNA binders attach through three cholesterol moieties at their base. Fluorescently labeled DNA binders and protein segregation or enrichment are visualized using fluorescence microscopy. Fluorescence intensity is measured through GUV contact linescans at both interface (1) and membrane exterior (2). The GUV Enrichment Index (GUV-EI) at the interface is calculated as the ratio of interface to exterior fluorescence. **(C)** GUV adhesion with short origami binder pair shows increased fluorescence at the interface. Enrichment index confirms short binder enrichment (GUV-EI > 1), indicating successful binding. Scale bar: 10 µm. Error bars: SD, N=26 vesicles pairs. **(D)** GUV contact with short binder pair and Fib5L showed no adhesion patches, with both components evenly distributed (GUV-EI ≈ 1). Scale bar: 10 µm. Error bars: SD, N=6 vesicle pairs. **(E)** With Fib5L bystander present, GUVs containing both long and short binders formed adhesion patches with strong origami fluorescence at the interface. Both binders were enriched (GUV-EI > 1) while Fib5L was excluded (GUV-EI < 1), confirming successful bystander barrier overcome. Scale bar: 10 µm. Error bars: SD, N=10 vesicles pairs.

The bystanders consisted of the same Fib 5L proteins that were fluorescently labeled at the N-terminus and anchored to vesicles via C-terminal His tags that associate with membrane Ni-NTA lipids. The short and long binders consisted of complementary DNA strands of two different lengths, mimicking ligand-receptor interaction through hybridization. These binders are anchored to GUVs via tetrahedral DNA origami, with three vertices carrying cholesterol anchors that insert into the bilayer, allowing stable anchoring of the binders on the vesicles. Each pair of binders contains a fluorescent marker for imaging. The long binder pairs have a maximum extension of approximately 20 nm and the short ones approximately 10 nm [37].

To quantify the distribution of molecules at interfaces, we measure fluorescence intensity across GUV pairs and calculate a GUV Enrichment Index (GUV-EI). This index is the ratio between the intensity at the interface (1) and the intensity at the free surface of vesicles (2) (Figure 4B). A GUV-EI > 1 indicates enrichment of molecules at the interface reflecting engagement of binders, a GUV-EI < 1 indicates their exclusion, and a GUV-EI ≈ 1 corresponds to a homogeneous distribution. When only short binder pairs decorate the surface of GUVs, we observe an increase in fluorescence at the interface (GUV-EI = 1.7 ±0.4) (Figure 4C), confirming effective engagement of complementary short binders.

However, when we add the bystander protein to the surface of one of the vesicles, we no longer observe enrichment of short binders at the contact point between vesicle pairs (Figure 4D), resulting in an enrichment index close to 1 (GUV-EI = 0.97 ±0.07). The Fib5L bystander proteins also remain homogeneously distributed (GUV-EI = 0.96 ±0.11), with neither enrichment nor depletion at the contact point. This demonstrates the blocking role of Fib5L, consistent with its height (∼17 nm), which is too large to allow engagement of the short binders.

We then tested whether adding long binders to the system (∼20 nm) would overcome the Fib5L bystander barrier and promote short binder engagement, as we had observed in the case of macrophages and target particles decorated with P-selectin (Figure 3). Consistent with the macrophage data, we observe a recovery of adhesion between vesicle pairs in the presence of long DNA binders, with enrichment of the short binders at the contact (GUV-EI = 1.4 ±0.6) and exclusion of the bystanders (GUV-EI = 0.3 ±0.2) (Figure 4E). We found that the long binders are also enriched at the interface (GUV-EI = 2.7±1), which may be due to the highly flexible single-stranded DNA region of the long binder that experiences a reduced entropic penalty compared to a tall binding protein and could remain at the membrane interface (Figure 4A). This observation of bystander protein exclusion and short binder engagement in even a simplified membrane interface with synthetic binders suggests that equilibrium thermodynamics may be sufficient to explain the observed kinetic enhancement at membrane interfaces.

### Theoretical modeling shows that energetically costly membrane deformations are required for short binder engagement in the presence of bystander proteins, slowing synapse formation

To better understand how bystanders and tall adhesive proteins influence close contact formation sustained by short binders like FcγR-IgG, we developed microscopic models that capture their fluctuating local concentrations and the varying shapes of their bound membranes. For short binders to engage, taller bystander molecules must be excluded from the contact zone. If the macrophage membrane were rigid, this would require complete removal of bystanders from the cell-target contact region (𝐴 = 1 𝜇𝑚^2^); an unrealistic scenario given the prohibitively high activation energy barrier (𝐹^‡^ = 200 𝑘*_B_*𝑇) for even a low surface density of bystanders (𝜌*_bystander_* = 200 𝜇𝑚^-2^).

Thus, we propose that fluctuating local abundances of proteins and membrane flexibility are key features of phagocytic synapse and cell-cell contacts. Specifically, we present a simple physical scenario involving a transition state that requires deformation of the macrophage membrane—formation of a dimple—allowing short binder engagement to nucleate within a small area with low local abundance of bystander molecules (Figure 5A). This bypasses the large energetic barrier of total bystander exclusion from the entire macrophage-target contact region prior to establishing strong favorable interactions between short binders, as also proposed for c-SMAC [38].

**Figure 5:**
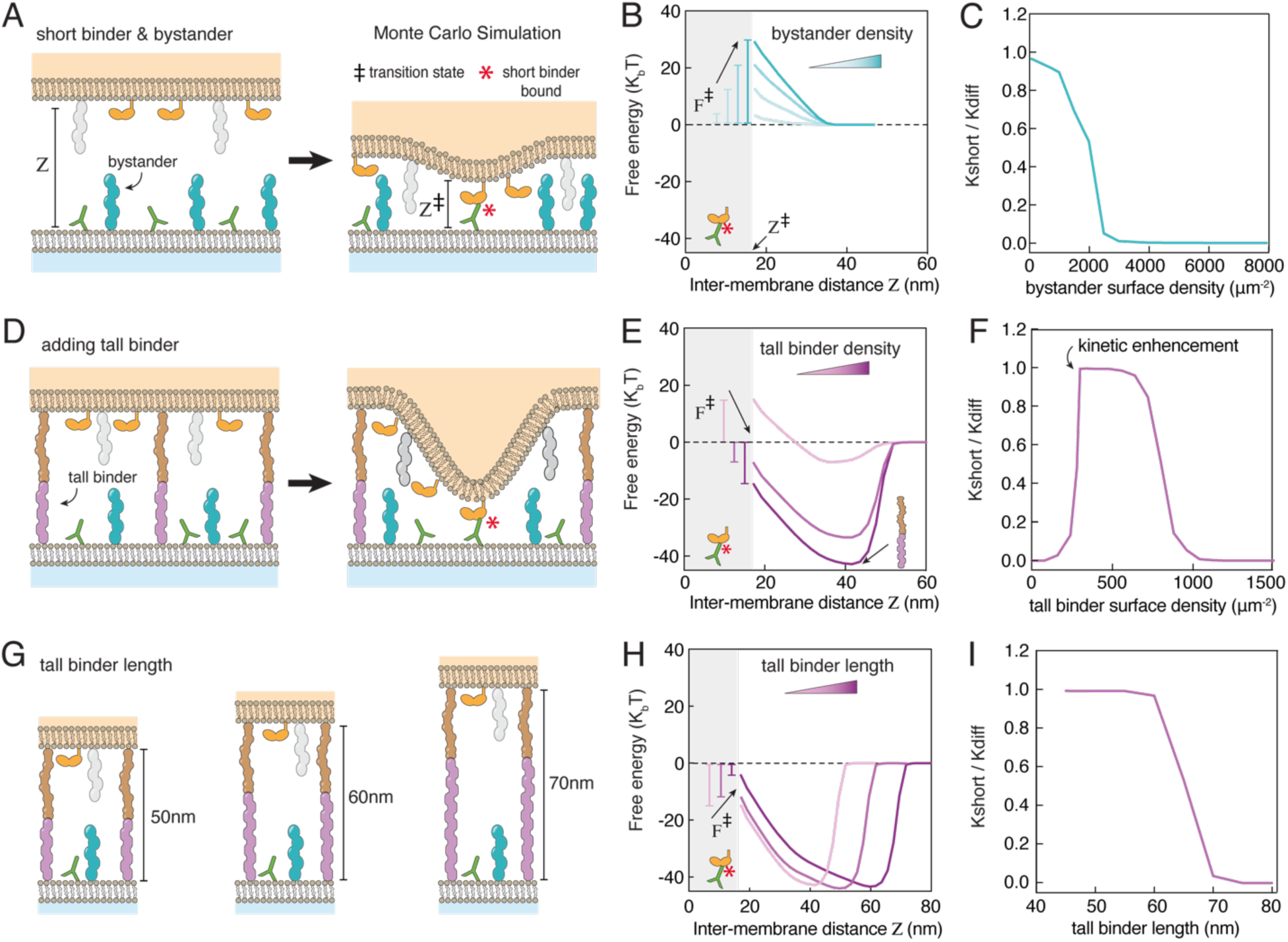
Monte Carlo simulations illustrate how bystander molecules block short binder engagement at a model macrophage-target interface and demonstrate that tall binders can overcome this barrier at specific densities and lengths. **(A)** Proposed mechanism shows how short proteins bind between a macrophage and target surface by displacing taller bystander proteins from the contact zone thanks to membrane deformation. The transition state exhibits a dimple-like distortion that brings the two membranes locally within the reach *Z*^‡^ of the short binder reducing thermodynamic cost of bystander exclusion. **(B)** Monte Carlo (MC) simulation of the interface enables calculation of the system’s free energy and confirms that increasing bystander molecule concentration raises the transition state free energy (*F*^‡^), impeding the engagement of short binders. **(C)** Increasing the bystander density raises the energetic barrier to short binder engagement, as shown by reduced kinetic rate of short binder engagement (*K_short_*) compared to the diffusion-limited rate constant in bystander-free conditions (*K_diff_*). **(D)** Tall binders that favorably bind across the membrane gap alter this pathway by stabilizing an intermediate membrane separation state. A larger dimple is required for short binder engagement and excluding both bystanders and tall proteins. **(E)** MC simulations of the interface reveal that the presence of tall binders (tall binder densities 80 µm^-2^, 320 µm^-2^ and 400 µm^-2^) creates an intermediate binding state that promote short binder binder engagement. **(F)** Beyond a critical concentration of tall binder, the engagement of short binders is kinetically enhanced until nearly reaching diffusion-limited rates. However, if the tall binder concentration is too high, excessive stabilization of the intermediate binding state blocks synapse formation kinetics. **(G)** Longer tall binders increase the inter-membrane distance of the intermediate state. **(H)** This raises the transition state free energy (*F*^‡^), as they require more extensive membrane deformation for short binder engagement. **(I)** The corresponding engagement rate constant shows that synapse formation is not kinetically enhanced for tall binders of sizes above than 60 nm. For parameters used in the MC simulations see SI Table S1.

For an interface containing only short binder pairs and bystanders, the energy required for dimple formation 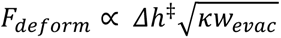 depends on three factors: the deformation’s vertical scale 𝛥ℎ^‡^, the membrane’s bending rigidity 𝜅, and the reversible work per unit area 𝑤*_evac_* = 𝜌*_bystander_*𝑘*_B_*𝑇 needed to evacuate bystanders from a membrane region (see SI text, Section 1). From this theory, we can draw several conclusions about the activation energy: it increases with (i) higher bystander density, (ii) greater deformation vertical scale, and (iii) increased membrane bending rigidity.

To further verify the basic features of these estimates, we performed Monte Carlo simulations of a microscopic model of macrophage and target membranes that includes short binders and bystander molecules (see SI text, Section 2). From these simulations, we computed the energy profile of interface formation as a function of *r*, the intermembrane distance, for different densities of bystander molecules. We found that the free energy barrier 𝐹^‡^ indeed increases as the bystander density increases (Figure 5B). The increase in this energy barrier causes a kinetic slowdown in short binder engagement (Figure 5C), which is consistent with the experimental results in Figure 1.

### Monte Carlo simulations suggest that specific densities of tall ligands help overcome barriers created by bystander molecules

We then investigated the effect of adding tall binder molecules on the kinetics of short binder engagement. We consider a population of tall binders (like the PSEL-PGSL1 pair) that can bind across the macrophage-target interface with sufficient strength to generate a metastable intermediate state at a membrane separation greater than the bystander’s height (Figure 5D). Introducing such an intermediate, with free energy 𝐹*_int_* < 0 and without lowering the maximum free energy 𝐹^‡^ along a barrier-crossing pathway, generically slows barrier-traversing kinetics. To enhance the kinetics of short binder engagement as observed in experiments, tall binders must act to stabilize the transition state. The schematic progression in Figure 5D illustrates how adhesion of a tall binder could do so. Here, membrane deformation begins from the intermediate membrane separation *h*, requiring a reversible work 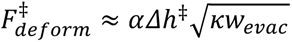 to initiate receptor engagement. The modified deformation scale 𝛥ℎ^‡^ and evacuation free energy per unit area 𝑤*_evac_* are both *larger* than the corresponding values for bystander proteins alone, so that the cost of deformation is actually magnified by the presence of tall binders. Importantly, however, the free energy maximum 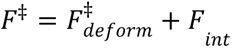, can be lower than in the absence of tall binders (see SI text, Section 1.B).

To study this effect, we performed a Monte Carlo simulation of membrane interfaces in the presence of tall ligands. The corresponding free energy profiles as a function of intermembrane space *r* show an intermediate minimum near the height of the tall ligand complex (Figure 5E). Increasing the tall ligand density lowers the free energy of the intermediate state and decreases the free energy barrier 𝐹^‡^. The kinetic rate of short binder engagement (*K_short_*) shows three distinct regimes as the density of long binders increases (Figure 5F). At low tall binder concentrations, stable intermediate states cannot form, so long binders have no effect on short binder engagement kinetic rate. Above a certain density threshold, tall ligand binding kinetically enhances synapse formation to reach uninhibited kinetic rates. This enhancement plateaus as concentrations become too high, at which point the intermediate state becomes overly stable, resulting in a collapse of the kinetic rates. The extent of the kinetically enhanced regime can be further modulated by varying the relative density of the tall binder pairs on both macrophage and target membranes (Figure S4). The three distinct regimes as a function of tall binder density observed in the simulations align with our experimental results showing that the phagocytic influence of P-selectin density is non-monotonic and has an optimum (Figure 3).

Finally, we investigated how changing the length of tall binders would affect the transition state free energy and subsequent kinetic rate of short binder engagement. We performed Monte Carlo simulation with tall binders of different lengths by varying the height of the tall binder on the target membrane, keeping all other molecular heights constant, and computed the resulting free energy profiles (Figure 5G,H). With increasing length of tall binders, more severe membrane deformation is required to reach the transition state, yielding higher free energy barriers 𝐹^‡^ but only slight changes in stability of the intermediate state. This results in a reduction of the kinetic rate of short binder engagement (Figure 5I), indicating that there may be an optimal height for the tall binders that is only slightly taller than the height of the bystander proteins.

## Discussion

The crowded environment of cell surfaces can hamper the formation of close cell-cell contacts. This is a particular challenge for macrophages, where antibodies targeting short antigens (<10 nm) are known to promote phagocytosis better than those against tall antigens (>10 nm) because close membrane contact is needed to exclude the inhibitory phosphatase CD45 and trigger phagocytosis [8]. How can short Fcγ receptors on macrophages overcome surface crowding and reach antibodies on short antigens that are hidden by surrounding taller bystander proteins? Here we demonstrate experimentally, theoretically, and with simulations that tall adhesive proteins at macrophage phagocytic synapses can promote antibody binding in the presence of bystander molecules that would otherwise inhibit phagocytosis.

Starting with antibody-coated target particles that are robustly phagocytosed, we show that addition of non-binding fibronectin domain repeat proteins can reduce phagocytosis by blocking Fcγ receptor engagement. Imaging of interface formation on supported lipid bilayers confirms that IgG-FcγR clustering is reduced and slowed as bystander protein density and height is increased. Surprisingly, we find that addition of another tall protein to the target particle—P-selectin, which binds the tall receptor PSGL-1 on macrophages—can restore phagocytic efficiency at low surface densities. This effect is only observed for a limited range of low P-selectin surface densities, as higher surface densities block phagocytosis. This suggests that while the presence of additional tall adhesive proteins adds to the crowding energy barrier preventing close contact, there is a regime of tall binder surface densities that can enhance the kinetics of IgG-FcγR binding and subsequent phagocytosis. To further test this idea, we reconstituted membrane interface formation in the presence of bystander proteins using GUVs together with DNA-based short and long binders. Consistent with our macrophage findings, we observed that tall binders could restore bystander protein exclusion and short binder engagement.

Theory and computer simulations suggest a generic physical mechanism for the observed kinetic enhancement of close membrane contact, in which long adhesive molecules mitigate the reduced kinetics of short binder engagement caused by taller bystander molecules. Simply stabilizing an intermediate state where the membranes are held together—but not in close contact—is not sufficient to explain the observed change in kinetics. While this might increase the chances of surface encounters, the process would remain inefficient since the system must still overcome a barrier to escape this intermediate state and achieve close contact. We argue that the intermediate state must involve deforming macrophage and/or target surfaces, allowing short adhesive molecules binding to occur while gradually excluding bystander proteins. Adding tall binders actually increases the cost of this deformation, but the transition state retains a large number of adhesive contacts.

We show that a net decrease in energy barrier height due to tall binders raises the rate of short binder engagement for low concentration and/or binding strength of the tall binders. However, with extensive and/or strong adhesion, escaping the proximal intermediate state becomes the limiting factor. In this regime tall binders instead inhibit the kinetics of short binder engagement. In the case of macrophages, since the intermediate state stability increases with the cell’s large footprint, the shift from enhancement to inhibition occurs at relatively low levels of adhesion, consistent with our experimental measurements. Our computational model was developed to validate the mechanistic principles observed in macrophage phagocytosis. While the model employs simplifying assumptions, it nevertheless successfully captured the experimental trends observed in our macrophage experiments. A more comprehensive model would account for additional features such as flexibility of binding molecules, which is likely to affect the energetics of the intermediate state. We suspect that binders with different measurable flexibilities [39] could lead to different kinetic enhancement.

Given that crowded membranes and multiple potential binders are often present at cell-cell interfaces, bystander-mediated blocking and kinetic enhancement by tall binders may be under-appreciated phenomena that occur in many natural systems. Efficient T-cell activation is known to require the binding of multiple co-receptors in addition to the TCR, including CD2, CD28, and the relatively tall integrin LFA-1, which binds to ICAM-1 on the target cell [40,41]. This leads us to speculate that perhaps a secondary role of some co-receptors is to provide kinetic enhancement of TCR interface formation, like the kinetic enhancement provided by P-selectin in our experiments on macrophage ADCP. Similarly, antibody dependent cellular cytotoxicity (ADCC) by Natural Killer cells, known to be an important contributor to cancer cell clearance during immunotherapy [42], relies on FcγR binding to IgG opsonized antigens on the target cell surface and is likely affected by bystander proteins.

Our observation of kinetic enhancement of ADCP mediated by tall binding proteins could potentially be utilized to improve existing antibody-based immunotherapies. In cases where an antibody is targeted to a short tumor antigen, kinetic enhancement can be achieved by forming a temporary interface between long molecules. Molecules with the ability to bind surface proteins on two different cells, as has been previously reported with bispecific antibodies [43] and bispecific T-cell Engagers (BiTEs) [44], could be used in combination with traditional immunotherapeutic antibodies that trigger ADCP and ADCC. Perhaps a ‘Big-BiTE’ therapeutic designed to form a tall interface with a gap just larger than the average height of bystander proteins could help to promote short immune receptor engagement. This would be particularly helpful for improving the efficacy of antibodies targeting tumor-associated short antigens close to the plasma membrane. For example, CD20, a B-cell associated antigen that is expressed on both healthy and malignant cells, has been successfully targeted by anti-CD20 IgG to clear B-cell lymphomas [45] However, CD20 is a very short antigen with multiple transmembrane domains and two short extracellular loops that can act as epitopes [46]. This fact, combined with evidence that anti-CD20 induced B-cell clearance relies on ADCP and ADCC by macrophages and Natural Killer cells [47], makes CD20 a prime target for a therapy that could be improved upon by kinetic enhancement.

We expect the need to mitigate kinetic frustrations imposed by bystander proteins to arise in many settings. Re-engineering transition states with long adhesive molecules is a simple strategy for doing so. Given the sensitivities to adhesion strength and distance we have observed, one can imagine not only rescuing efficient receptor engagement but also modulating it precisely in space and time, efficiently discriminating among receptor pairs according to their height. Realizing such control in engineered cell systems, and exploring the implications in natural biological contexts, are exciting directions for future work.

## Acknowledgments

We would like to thank Fletcher laboratory members and Geissler group members for their input on this project. Thanks to Dr. Siddhansh Agarwal for his contributions to the theory section. This work was supported by NIH R01 GM134137 and the NSF Center for Cellular Construction (DBI-1548297) to D.A.F. A.C. was funded by an EMBO Postdoctoral Fellowship for this work. D.A.F. is a Chan Zuckerberg Biohub Investigator. In memory of Phill Geissler.

## Material and Methods

### RAW264.7 cell culture

RAW264.7 macrophage-like cells were obtained from the UC Berkeley Cell Culture Facility. The cells were cultured in RPMI media (Corning) with supplemental Penicillin-Streptomycin (1%) and Fetal Bovine Serum (10%). Cells were passaged at a ratio of 1:5 every two days. After 2 months of passaging, they were replaced with a new batch of low passage number cells to prevent phenotypic drift from the normal cell line state.

### Assaying phagocytosis

All phagocytosis assays were performed in accordance with the steps outlined in our previously published protocol [19]. Briefly, reconstituted target particles were made by coating 4 µm diameter glass beads (Bangs Laboratories) with a fluid lipid bilayer. His-tagged proteins were coupled to the bilayer via interaction with Ni-NTA lipids in the bilayer. Anti-biotin IgG (eBioscience, clone BK-1/39, Alexa Fluor 488 label) was targeted to biotinylated lipids in the bilayer. Reconstituted target particles were incubated with RAW264.7 macrophage-like cells in a 96-well plate for 30 minutes at 37 °C after which excess beads were washed from the wells and CellTracker Green CMFDA dye (Invitrogen) was added to the wells to fluorescently label the cells. Phagocytosis was quantified by capturing fluorescence images of cells and targets, images which were then analyzed using an automated CellProfiler (Broad Institute) analysis pipeline. The phagocytic index for each condition was determined by measuring the average internalized target particle fluorescence intensity per cell. All experiments were repeated at least 3 times on 3 separate days. To obtain final phagocytic indices with errors, each daily phagocytic index was normalized to the average phagocytic index across all conditions to obtain a normalized phagocytic index that accounts for day-to-day global variations in phagocytosis. Error bars represent standard error across all replicates of a given condition.

### Reconstitution of target particles

Small Unilamellar Vesicles (SUVs) were made by mixing lipids dissolved in chloroform, followed by desiccation, rehydration and sonication. Lipid bilayers coating reconstituted target particles contained the following: 0.5% Biotin PE (Avanti), 0.4% 647 DOPE (Atto-Tec), 0.5%, 1.6%, 2.8% or 5% Ni-NTA (Avanti), and the remainder POPC (Avanti). For measurements of Fibcon densities, 647 DOPE was left out so that it would not interfere with the fluorescence signal of the fluorescently labeled Fibcon proteins. During protein conjugation steps, Fibcon proteins were incubated with lipid coated target particles at a concentration of 50 nM and anti-biotin IgG at a concentration at 125 ng/mL. For experiments in which P-selectin was displayed on target surfaces alongside Fibcon proteins and anti-biotin IgG, His-tagged P-selectin (Sino Biological) and Fibcon were added at different ratios, while keeping the total protein concentration fixed at 50 nM.

### Fibcon family protein expression and purification

Fibcon proteins (Fib1L, Fib3L, Fib5L) were expressed in Rosetta DE3 competent *E. coli* cells (EMD Millipore). After transfection with the plasmid, cells were grown at 37 °C until they reached an OD of 0.8, at which point they were induced with 0.3 mM IPTG and incubated overnight at 18 °C. Cells were harvested and resuspended in PBS with 0.5 mM TCEP and 10 mM imidazole followed by cell lysis via sonication. The lysate was clarified by centrifugation at 20,000g for 45 minutes. Protein was ultimately purified by loading onto a His-Trap HP column (GE Healthcare) and eluting with an imidazole gradient on an AKTA Pure (GE Healthcare) chromatography system. Protein peak fractions were further purified and buffer exchanged into PBS via gel-chromatography on a Superdex 200 column (GE Healthcare). TCEP (0.5 mM) was added to each protein solution to ensure reduction of disulfide bonds and proteins were flash frozen and stored in small aliquots.

### Measuring protein densities on target particles

Fib3L proteins were labeled with Alexa Fluor 647 at a single C-terminal cysteine labeling site using Alexa Fluor 647 C_2_ Maleimide (Invitrogen). Reconstituted target particles were created using the same lipid compositions as used in the phagocytosis assays, but without any fluorescent 647 DOPE lipid. Just as in the phagocytosis assays, targets with different fractions of Ni-NTA lipids were used to display different densities of Fibcon proteins. Specifically, targets with Ni-NTA fractions of 0.5%, 1.6%, 2.8% and 5% were made. Fluorescent Fib3L proteins were used to coat the particles at the typical concentration of 50 nM along with 125 ng/mL anti-biotin IgG. These target particles were analyzed for Fib3L fluorescence using flow cytometry and the average fluorescence for each condition was recorded with at least 10,000 particles measured per condition. Using the same instrument settings, calibrated Alexa Fluor 647 MESF beads (Bangs Laboratories) were measured to create a conversion between fluorescence intensity signal and absolute fluorophore counts (Figure S1). Fluorophore densities were determined by taking the total number of fluorophores measured on each condition of particles and dividing by the surface area of the 4 µm beads. The fluorophore density was considered equal to the Fibcon protein density because each Fibcon protein was labeled with a single fluorophore.

### Preparation of Supported Lipid Bilayers (SLBs)

Planar supported lipid bilayers (SLBs) were prepared analogously to reconstituted target particles by incubating 50 uL SUV solution and 50 uL MOPS buffer (25 mM, MOPS, 125 mM NaCl, pH 7.4) in a PDMS chamber with RCA cleaned glass coverslips. After 20 minutes, the well was washed 5x with PBS to wash away excess SUVs while ensuring that the chamber does not dry out. Fibcon proteins and anti-biotin IgG were added to the well at respective concentrations of 50 nM and 125 ng/mL and allowed to incubate for 20 minutes. Finally, the well was washed 5x with PBS to remove excess Fibcon proteins and IgG.

### TIRF imaging

Live cell imaging was performed using a stage-top incubation chamber (Okolab) to ensure 37 °C temperatures and 5% CO_2_. For time-lapse image acquisitions macrophages were dropped onto SLBs inside the incubation chamber and immediately imaged. Micro-manager software was used to automatically control the microscope (Nikon) and capture both TIRF and RICM images every 30 seconds for multiple fields of view. For endpoint image acquisitions macrophages were incubated on SLBs in the stage-top incubator for 30 minutes prior to imaging.

### Enrichment analysis

Images collected from TIRF imaging were analyzed using Fiji analysis software. Individual cells were identified in the RICM channel of each image. A mask around the boundary of each cell was drawn and the mask was applied to the corresponding TIRF image. The average fluorescence intensity emanating from Alexa Fluor 488 labeled anti-biotin IgG within each cell mask was measured and the enrichment index (EI) was determined by dividing each of these values by the average background fluorescence intensity of the same field of view. Each of these experiments were repeated at least 3 times on 3 separate days. Data points from all individual experiments were combined for each condition to comprise the final data.

### Assaying target particle binding

Reconstituted target particles were made in the same way described above (see *“Reconstitution of target particles”*) and in [19]. RAW264.7 cells were labeled with CellTracker Green CMFDA and washed 2x with PBS followed by final resuspension in cell media. Cells were diluted to a concentration of 350 cells/µL and 100 µL of cells were added to each tube. 100 µL of target particle suspension containing ∼500,000 particles was added to each tube and the tubes were incubated for 30 minutes at 4 °C for to prevent phagocytosis and under constant rotation to prevent settling. After 30 minutes, each tube was run through a flow cytometer and fluorescence data for at least 5,000 macrophages were collected. All tubes were kept on ice prior to loading into the flow cytometer and the entire run time was kept under 30 seconds to ensure that bound particles were not phagocytosed as the sample warmed to room temperature. A low flow rate was used to prevent unnecessary shearing of bound targets from cells. Results were quantified by plotting particle fluorescence (647 DOPE) vs. macrophage fluorescence (CMFDA). The fraction of macrophages with targets bound to their surface was determined by dividing the number of macrophages with targets bound to their surface (647 DOPE+, CMFDA+) by the number of all macrophages (CMFDA+).

### Giant Unilamellar Vesicles (GUVs) formation

For all experiments, we used a consistent lipid mixture to form GUV membranes (POPC 74.5%; Peg2K 0.5%; Chol 20%, DSG-NTA(Ni) 5%) when the blocker Fib5L should be anchored to the GUVs, and (POPC 79.5%; Peg2K 0.5%; Chol 20%) otherwise. To prepare GUVs, we spread 0.25 mg of the lipid mixture uniformly on indium tin oxide (ITO)-coated slides. After vacuum drying for greater than 30 minutes to remove chloroform, we assembled a capacitor using two lipid-coated slides as electrodes separated by a 0.3-mm rubber isolator septum. We filled the chamber with ∼200 μL of 350 mM sucrose solution (∼350 mOsm). GUVs (10–100 μm diameter) were formed by electroformation, applying 1.5 V AC at 10 Hz for 1 hour to the capacitor, following established protocols [48]

### Formation of DNA tetrahedra (short and long binder)

To assemble the DNA tetrahedron binder, we combined 0.5 µL of each oligonucleotide building block (Tet-B, Tet-C, Tet-D, and the variable strand) in PCR tubes and added 48 µL of PBS 1X before vortexing and placing the mixed solution in a thermal cycler. The thermal cycling program ran for 90 minutes through the following steps: initial hold at 20°C, heating to 90°C and hold for 10 min, cooling at -1.5°C/minute until reaching 20°C, and finally holding at 4°C. The final concentration of the origami tetrahedron in the solution is 1 µM. We characterize the formation of origami structure using 5% TBE gel (Figure S3). The binding domain of the synthetic origami binder has the same length (12 nucleotides) for both tall and short variants, but the short binder is designed with a stronger binding domain, having 80% GC content compared to the tall binder’s 25% GC content. See Supplementary Table 2 for the complete sequences used to assemble the origami binders.

### GUV functionalization with short and long DNA binders and/or Fib5L blocker molecules

The origami stock solutions are diluted in PBS 1X to obtain 100 µL of 500 nM solution in an Eppendorf tube containing 50 nM of Fib5L blockers. 5 µL of GUV solution (with 5% nickel lipids when Fib5L blocker needs to be anchored) are added to this solution, gently homogenized using a micropipette, then incubated on the rotisserie for 5 minutes. To eliminate excess origami (and/or Fib5L), the tube is placed vertically to allow GUV sedimentation. Then, 10 µL of supernatant are collected and mixed with 90 µL of PBS; this washing step is repeated twice. The purified solution is maintained on the rotisserie to prevent sedimentation of functionalized GUVs until use.

### Formation of GUV-GUV pairs

Once the surface of both complementary GUV types is decorated with the necessary origami and blockers, they are brought into contact by mixing 5 µL of each complementary GUV solution in 100 µL of PBS. The mixture is deposited on a coverslip previously passivated by incubation for 1 minute with 10% BSA, then rinsed 3 times with buffer (PBS), to prevent vesicle rupture.

### Imaging and measurement of enrichment index at GUV-GUV interface

The formed GUV-GUV pairs are then imaged at their equatorial plane on a Nikon Ti-2 microscope equipped with a spinning disk (Yokogawa CSU-X1) through a SR plan Apo IR 60X/1.27 WI objective. The fluorescence intensity of the different molecules present (DNA origami binder and Fib5L blockers) is measured by a linescan and allows calculation of an enrichment index at the contact between vesicles (GUV-EI) by taking the ratio of fluorescence intensity at the contact normalized by that outside the contact [49].

### Theory overview

The theoretical framework developed in this work models the interaction between macrophages and target particles, focusing on key “dimple” formation in the macrophage membrane during receptor engagement. The model quantifies the energetic cost of membrane deformation using Helfrich’s small-gradient approximation, accounting for both elastic energy and the thermodynamic cost of bystander protein evacuation from the contact zone. The kinetics of receptor engagement are described using overdamped Langevin dynamics, with separation vector r between a macrophage and a target particle treated as reaction coordinate. The steady-state solution to the resulting Smoluchowski equation, with appropriate boundary conditions, yields phagocytosis rates relative to the diffusion-limited case. This framework particularly elucidates how tall adhesive proteins can mitigate the kinetic effects of bystander molecules through barrier reduction. See SI text for full description of the theory framework.

### Monte Carlo simulations overview

Simulations were performed to investigate microscopic fluctuations in bystander-hindered phagocytosis, using a model adapted from [50]. The model represents the macrophage and target membrane topography on a discrete lattice, with membrane-bound proteins treated as a noninteracting lattice of rigid rod-like molecules. The system consists of two two-dimensional lattices representing the macrophage and target membranes, with surface anchored molecules – short binder and tall binder representing respectively FcγR/IgG or P-selectin/PSGL-1 and blocker molecules representing the macrophage and target bystander proteins – occupying sites subject to spatial and energetic constraints. The simulation employed a grand canonical ensemble approach, where protein population fluctuations were integrated out to yield a free energy function governing membrane separation. Changes in membrane height were proposed at random lattice sites and accepted according to the Metropolis criterion, accounting for membrane elastic energy, protein-mediated free energy changes, and a biasing potential. Free energy profiles were computed along a membrane gap coordinate (*Z*) using umbrella sampling. The simulation was performed on a 0.7 μm² elastic sheet discretized into a 70×70 grid of 12×12 nm² patches. The system was equilibrated for 2×10⁵ Monte Carlo sweeps, before data was collected from an additional number of MC sweeps between 8x10⁵and 2x10^6^. Refer to SI text Section 2 for a full description of the Monte Carlo simulations.

