## Supporting Information for "Overcoming steric inhibition of antibody-dependent phagocytosis with tall adhesions"

##### This PDF file includes:

Supporting text
Tables S1 to S2
Figs. S1 to S4

### Supporting Information Text

#### 1. Theoretical modeling

**A. Dimple Thermodynamics.** The transition state we propose for bystander-hindered receptor engagement involves a localized deformation of the macrophage membrane that we refer to as a “dimple”. Here we estimate the reversible work $F_{\mathrm{deform}}$ required to create such a dimple, beginning from an undeformed membrane at height $h_0$ that allows tall proteins to populate the entire interfacial contact zone. The transition state, sketched in Fig. T1, remains undeformed except in a lateral region of radius $R$. The tip of this circularly symmetric dimple extends a distance $\Delta h$ below $h_0$. We adopt a lateral coordinate system $x$ whose origin coincides with the dimple tip, so that the height profile $h(x)$ depends only on the scalar $x = |x|$. In these terms, the geometry described above is specified by the conditions $h(x) = h_0$ and $h(0) = h_0 - \Delta h$.

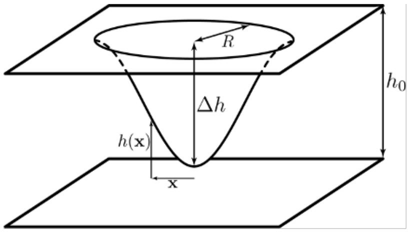


**Fig. T1. Dimple Thermodynamics Parameterization.**

The membrane’s elastic energy $E_{\mathrm{elastic}}$, in the small-gradient limit of Helfrich’s model, can be written

$$E_{\mathrm{elastic}} = \frac{1}{2}\kappa \int dx (\nabla^2 h)^2 \quad [1]$$

A finite elastic energy thus requires additional boundary conditions $h'(R) = h'(0) = 0$. Subject to these constraints, we utilize a smooth, one-parameter interpolation for the dimple profile where the functional $E_{\mathrm{elastic}}[h(x)]$ is minimized by a height profile

$$h(x) = h_0 - \Delta h \left[1 - \left(1+\sqrt{2}\right)\left(\frac{x}{R}\right)^2 + \sqrt{2}\left(\frac{x}{R}\right)^{2+\sqrt{2}}\right] \quad [2]$$

for which

$$E_{\mathrm{elastic}} = 4\pi\left(1+\sqrt{2}\right)\kappa\left(\frac{\Delta h}{R}\right)^2 \quad [3]$$

The region of the membrane where $h(x) < h_0$ incurs an additional thermodynamic cost $\pi R^2 w_{\mathrm{evac}}$, where $w_{\mathrm{evac}}$ is the reversible work per unit area required to evacuate proteins (and/or protein complexes) whose heights exceed the local membrane gap. In general, protrusion of the dimple would exclude progressively more protein species as $h(x)$ decreases; here, for simplicity, we consider only proteins/complexes of a single height $h_0$. For a target substrate displaying only IgG and Fibcons, $h_0$, represents the height of bystander proteins, and $w_{\mathrm{evac}} = \rho_{\mathrm{bystander}} k_B T$ originates in their translational entropy. For a tall adhesive protein like P-selectin, $w_{\mathrm{evac}}$ may instead be dominated by the disruption of favorable protein complexation across the membrane gap.

The total free energy of deformation, $E_{\mathrm{elastic}} + \pi R^2 w_{\mathrm{evac}}$, is minimum for a dimple width $R = \left(\frac{4(1+\sqrt{2})\kappa\Delta h^2}{w_{\mathrm{evac}}}\right)^{1/4}$, giving

$$F_{\mathrm{deform}} = 4\pi\Delta h\sqrt{(1+\sqrt{2})\kappa w_{\mathrm{evac}}} \quad [4]$$

**B. Steady state kinetics.** To predict the rate of membrane receptor engagement from computed free energy profiles $F(r)$, we consider an overdamped Langevin dynamics of the separation vector r between a macrophage and a target particle, with $F(r)$ as an effective potential energy. In doing so, we assume that all other degrees of freedom relax rapidly by comparison.

For a tagged macrophage/target pair, the probability distribution p(r,t) satisfies a Smoluchowski equation

$$\frac{\partial p}{\partial t} = -\nabla \cdot J \tag{5}$$

with flux $J = -De^{-\beta F}\nabla(e^{\beta F}p)$, where $\beta = (k_B T)^{-1}$ and $D$ is the diffusion coefficient for their relative motion. The rate of phagocytosis (here short binder engagement) is determined from a steady-state solution to Eq. 5, with boundary conditions describing (i) irreversible receptor engagement when the distance $r = |r|$ reaches a minimum value $r_0$ (i.e., $p(r_0) = 0$), and (ii) asymptotic approach to a constant probability $p$ at large $r$ (i.e., $p(r) \to p$ as $r \to \infty$).

We focus on the rate of short binder engagement $k_{\text{short}}$ relative to the diffusion-limited value $k_{\text{diff}}$ that would be obtained in the absence of a free energy barrier,

$$\kappa = \frac{k_{\text{short}}}{k_{\text{diff}}} = -\hat{r} \cdot J(r_0)\frac{r_0 V}{D} \tag{6}$$

where $\hat{r}$ is a radial unit vector and $V$ is the macroscopic sample volume. A standard Kramers analysis of this boundary value problem yields

$$\kappa = Vp(r)e^{\beta F(r)}\left(r_0\int_{r_0}^{r} dr\,\frac{e^{\beta F(r)}}{r^2}\right)^{-1} \tag{7}$$

for all $r$, and in particular

$$\kappa = \frac{Vp}{r_0}\left(1 + r_0\int_{r_0}^{r_{\text{contact}}} dr\,\frac{e^{\beta F(r)}}{r^2}\right)^{-1} \tag{8}$$

where $r_{\text{contact}}$ is a contact distance beyond which $F(r) = 0$. (See Fig. T2) Note that $r_0$ and $r_{\text{contact}}$ are both comparable to the macrophage's overall size (∼20 microns in diameter), while $r_{\text{contact}} - r_0$ is a membrane gap scale of roughly 10 nm.

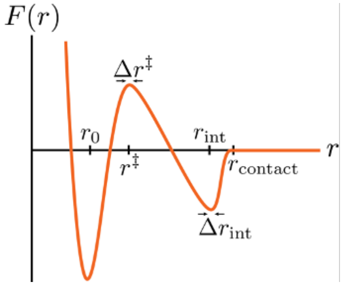


**Fig. T2. Free energy profile of membrane contacts.**

Kinetic suppression from a deeply metastable intermediate manifests in this description as a depletion of $p$, with substantial population sequestered in the distance interval $r_{\text{int}} \pm\Delta\ r_{\text{int}}$. Here we account for the depletion roughly, by imagining macrophages to be stationary, establishing $N_{\text{macro}}$ localized traps where a target particle tends to dwell on the macrophages surface at $r_0$. Normalization of the steady-state distribution $p(r)$ then gives

$$1 = N_{\text{macro}}\, r_0 \int_{r_0}^{r_{\text{contact}}} 4\pi r^2 p(r)\, dr + Vp \tag{9}$$

Combining Eqs. 7-9, we finally obtain

$$\kappa = (1 + b + c\rho_{macro} r_{contact}^3)^{-1} \quad [10]$$

where $\rho_{\text{macro}} = N_{\text{macro}}/V$.

$$b = r_0 \int_{r_0}^{r_{\text{contact}}} dr\, \frac{e^{\beta F(r)}}{r^2} \quad [11]$$

and

$$c = r_{\text{contact}}^{-2} \int_{r_0}^{r_{\text{contact}}} dr\, 4\pi r^2 e^{-\beta F(r)} \int_{r_0}^{r} dr\, \frac{e^{\beta F(r)}}{r^2} \quad [12]$$

Saddle point approximations for these integrals yield

$$k_{\text{short}} = \frac{k_{\text{diff}}}{\left(1 + \frac{\Delta r^{\ddagger}}{r_{\text{contact}}} e^{F^{\ddagger}/k_B T} + \alpha e^{(F^{\ddagger} - F_{\text{int}})/k_B T}\right)} \quad [13]$$

Here $k_{\text{diff}}$ is the rate of diffusion-limited short binder engagement, in which receptor engagement is not impeded by a free energy barrier, $\Delta r_{\text{int}}$ and $\Delta r^{\ddagger}$ are nm-scale membrane gap ranges corresponding to intermediate and transition states, and $r_{\text{contact}} \approx 1$µm is the distance between the center of a macrophage and the center of a target particle when they contact. The number density of macrophages $\rho_{\text{ macro}}$ is on the order of $10^{11}$ in our experiments, so $\alpha = 4\pi\rho_{\text{macro}} r_{\text{ contact}} \Delta r^{\ddagger} \Delta r_{\text{int}}$ is set to $1\text{x}10^{-11}$ to resemble experimental conditions. Eq. 13 makes clear that an intermediate state which does not lower $F^{\ddagger}$ cannot offer kinetic enhancement. For the free energy profiles we have computed, one term in the denominator of the right-hand side typically dominates over the others, establishing three distinct dynamical regimes: (i) Without a high barrier or a significantly stable intermediate ($F^{\ddagger}$, $F_{\text{int}} \approx 0$), kinetics is diffusion-limited. (ii) Absent a deeply metastable intermediate, a high barrier ($F^{\ddagger} \gg k_B T$) gives the simple transition state theory estimate $k_{\text{short}} \approx k_{\text{diff}}\; e^{-F^{\ddagger}/k_B T}$. In this regime tall adhesive proteins can substantially offset kinetic effects of bystanders by lowering $F^{\ddagger}$. (iii) An interfacial free energy minimum so deep that $F^{\ddagger} - F_{\text{int}}$ exceeds even $k_B T |\ln \alpha|$ will suppress barrier crossing, regardless of the value of $F^{\ddagger}$.

#### 2. Simulations

**A. Monte Carlo Simulations.** To examine and quantify the role of microscopic fluctuations in bystander-hindered phagocytosis, we performed Monte Carlo simulations of a model adapted from the work of Liposwky and coworkers [1,2]. It resolves topography of the macrophage membrane at a discrete lateral scale $\Delta x_{\text{membrane}}$ (the target membrane is taken to be completely rigid), and treats the populations and spatial arrangements of membrane-bound proteins as a (laterally) noninteracting lattice gas. Proteins can occupy sites on a pair of two-dimensional lattices, representing the macrophage (upper) and target (lower) membranes, subject to constraints that (i) no more than one protein can simultaneously occupy a given lattice site, and (ii) the local vertical separation between upper and lower membranes must exceed $h_{\text{upper}} + h_{\text{lower}}$ in order to accomadate proteins of height $h_{\text{upper}}$and $h_{\text{lower}}$ in the upper and lower membranes. When an appropriate pair of proteins (Fc$\gamma$R/IgG or P-selectin/PGSL-1) are apposed across the membrane gap and separated vertically by less than a threshold distance $l_{\text{bind}}$, their interaction contributes a favorable energy of binding $-\epsilon^{(bind)}$.

For a fixed membrane shape, protein population fluctuations can be integrated out exactly in a grand canonical ensemble, yielding a free energy

$$F[h(x)] = \frac{\kappa}{2} \sum_x \left(\nabla^2 h\right)^2 + \sum_x U(h(x)) \quad [14]$$

that governs fluctuations in local membrane separation $h(x)$. Here, $\Delta x_{\text{protein}}$ is the lateral footprint of a protein (which we take to be the same for all proteins), $\nabla^2$ is a discrete approximation to the 2-d Laplacian operator, and $U(h)$ is an effective potential due to the constraints described above. In terms of possible protein occupation states $i$ (which may include protein(s) in the upper and/or lower membranes,

$$U(h) = -k_B T \ln \left( \frac{\Delta x_{\text{protein}}}{\Delta x_{\text{membrane}}} \right)^2 \ln \left( \sum_i \theta(h - h_i) z_i e^{\beta \epsilon_i^{(\text{bind})}} \theta(h_i + l_{\text{bind}} - h) \right) \qquad [15]$$

where $h_i$ is the summed height of proteins present, $z_i$ is the product of their activities, $\epsilon^{(bind)}$ is their binding energy (which vanishes in all states except Fc $\gamma$ R/IgG and P-selectin/PSGL-1), and $\theta(y)$ is the Heaviside step function. The activity of a protein in this description, expressed in terms of its surface density $\rho$ is $\rho {\Delta x_{\text{protein}}}^2/(1 - \rho \Delta x_{\text{protein}})$.

Our MC simulation consists of attempts to change $h(x)$ at a single, randomly chosen lattice site $x$. The change in height is also chosen at random from the interval [-dh, dh]. The proposed new height for the chosen lattice site's is accepted with the standard Metropolis probability $[1, e^{-\beta(\Delta F_{\text{membrane}} + \Delta U + \Delta F_{\text{bias}})}$, where $\Delta F_{\text{membrane}}$ is the associated change in the elastic energy of the membrane, $\Delta U$ is the change in free energy due to the proteins, and $\Delta F_{\text{bias}}$ is the change in a biasing potential designed to improve sampling of important but unlikely configurations.

Free energy profiles $F(r)$ were computed along a membrane gap coordinate $r = \min_n h(x)$ that parametrizes progress through the barrier-crossing pathways of Fig. T2 and is well-suited to the kinetic description of Eq. 5. To obtain $F(r)$, we compute and integrate the mean force $\langle f(r) \rangle \approx 2 * k_{\text{bias}} \langle (r - r_0) \rangle$ from the average deviation of $r$ away from the the value favored by a stiff harmonic bias potential $F_{bias} = k_{\text{bias}}(r - r_0)^2$.

A 0.7 µm$^2$ elastic sheet was discretized into a 70x70 square grid composed of 12x12 nm$^2$ patches. The values of the physical parameters used in the simulation are given in the table below. An umbrella sampling simulation protocol for r sampling was performed in a range of 15-80 nm. Each simulation was equilibrated for 2x10$^5$ Monte Carlo sweeps before data was collected from an additional number of MC sweeps between 8x10$^5$ and 2x10$^6$. Changes in height were drawn from the interval [-1.2, 1.2] nm, leading to a 60% acceptance rate.

**B. Monte Carlo Parameters.** Supporting Information Table 1 contains parameters used to perform Monte Carlo simulations of cell-target contacts.

| | Parameter | Value |
|---|---|---|
| 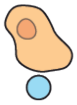 | Cell membrane bending modulus | 20 $k_B T$ |
| | Target (Bead) supported bilayer | rigid |
| 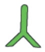 | Target Short binder (mimic IgG) | |
| | Length | 7.5 nm |
| | Density (molecules per μm²) | 200 /$\mu m^2$ |
| 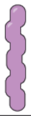 | Target tall binder (mimic P-Selectin) | |
| | Length | 30.0 nm |
| | *Density | 0 - 1600 /$\mu m^2$ |
| 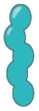 | Target Bystander (mimic FIB5L) | |
| | Length | 17.5 nm |
| | *Density | 8000 /$\mu m^2$ – PSEL density |
| | *Density Bystander Only Case | 50 - 10000 /$\mu m^2$ |
| 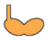 | Cell short binder (mimic FcR) | |
| | Length | 3.5 nm |
| | Density | 200 /$\mu m^2$ |
| 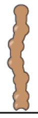 | Cell tall binder (mimic PSGL) | |
| | *Length | 15 - 50 nm |
| | Density | 200 /$\mu m^2$ |
| 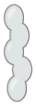 | Cell Bystander | |
| | Length | 17.5 nm |
| | Density | 50 - 10000 /$\mu m^2$ |
| | Cell + Target contact area | 0.7 $\mu m^2$ |
| | Molecular diameter | 4 nm |
| | Note: Quantities labeled with * are the sampling parameters | |

**Table S1. Monte Carlo Simulation Parameters.**

#### 3. DNA Oligonucleotides

**A. DNA Sequences.** Supporting Information Table 2 contains DNA sequences used to assemble DNA binding pairs.

| Names | Sequences | Description |
|---|---|---|
| Tet-B | CCGCGGTTGCAGCGCTCACGGCGTT/3CholTEG/ | Invariable strand to assemble tetrahedron |
| Tet-C | ACCGCGGTGGTGGGCTCGGAGCCTT/3CholTEG/ | Invariable strand to assemble tetrahedron |
| Tet-D | GGTCCGCTGCGCTGCTGGCTCCGTT/3CholTEG/ | Invariable strand to assemble tetrahedron |
| Short Binder | /5RhoR-XN/TTTTTCGGCCCGTGCTGTTGCGGACCTGCCCACCTCGCCGTG | Variable sequence to form binding pairs |
| Short Binder Complement | CAGCACGGGCCGTTGCGGACCTGCCCACCTCGCCGTG | Variable sequence to form binding pairs |
| Long Binder | /5Alex488N/TTTTTATCATATGACTTTTTTTTTTTTGCGGACCTGCCCACCTCGCCGTG | Variable sequence to form binding pairs |
| Long Binder Complement | AAGTCATATGATTTTTTTTTTTGCGGACCTGCCCACCTCGCCGTG | Variable sequence to form binding pairs |

**Table S2. DNA oligos containing modifications and sequences as ordered from IDT.**

#### 4. Supporting Figures

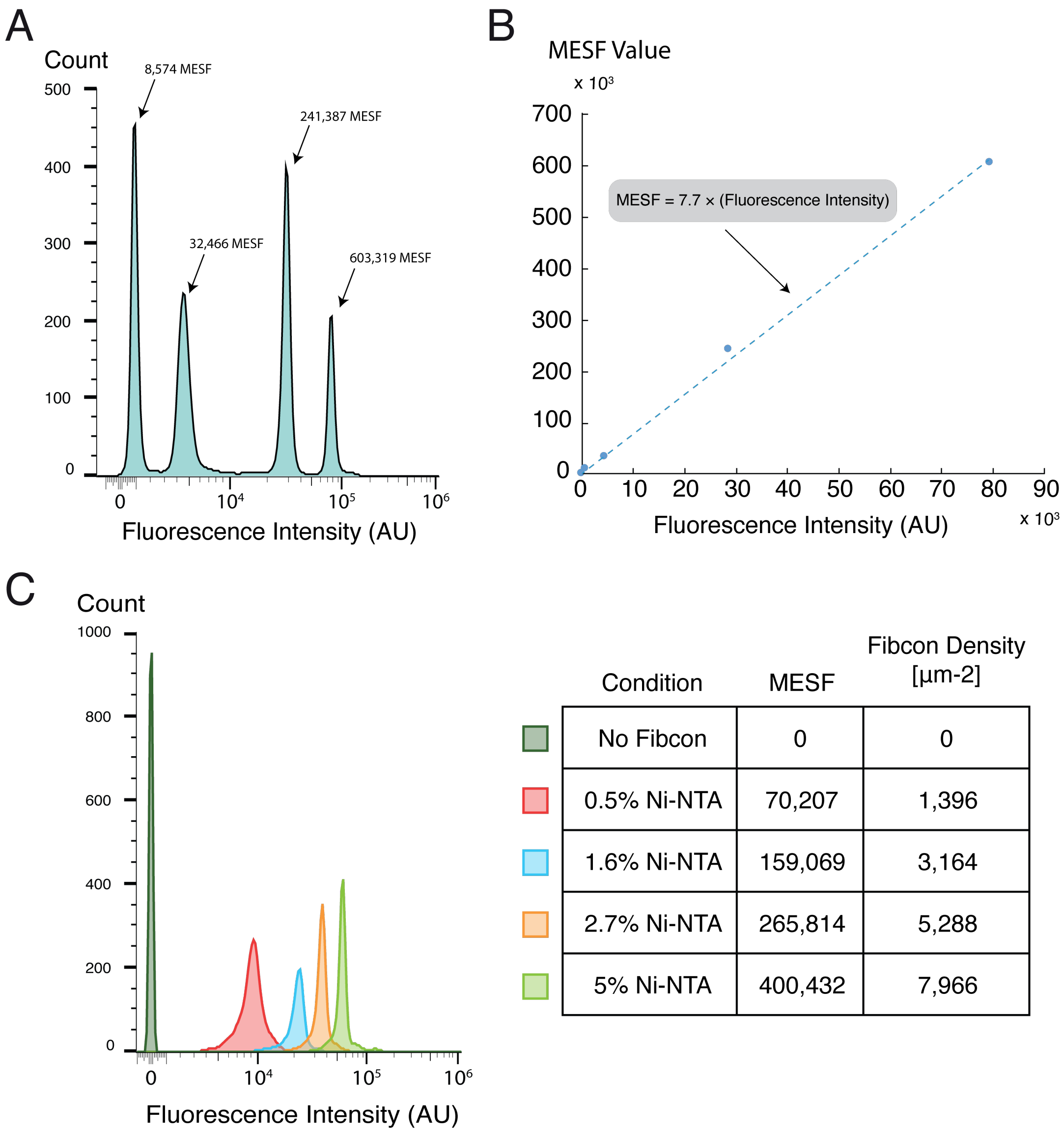


| Condition | MESF | Fibcon Density [µm-2] |
|---|---|---|
| No Fibcon | 0 | 0 |
| 0.5% Ni-NTA | 70,207 | 1,396 |
| 1.6% Ni-NTA | 159,069 | 3,164 |
| 2.7% Ni-NTA | 265,814 | 5,288 |
| 5% Ni-NTA | 400,432 | 7,966 |

**Fig. S1. Quantification of surface protein densities. A.** Histogram of measured fluorescence intensities for calibrated Alexa Fluor 647 MESF beads. **B.** Calibration curve of MESF values vs. fluorescence intensity for calibrated MESF beads. The fit line yields an equation for converting fluorescence intensity to MESF. **C.** Histogram of target particles displaying Fib3L that was fluorescently labeled with Alexa Fluor 647 using maleimide chemistry to react with a lone cysteine on the protein (yielding 1 fluorophore per Fib3L protein). Ni-NTA densities were varied to match those used in other experiments and the measured fluorescence intensity was converted to MESF and ultimately to an absolute Fibcon density by dividing by the surface area of the target (4 µm diameter).

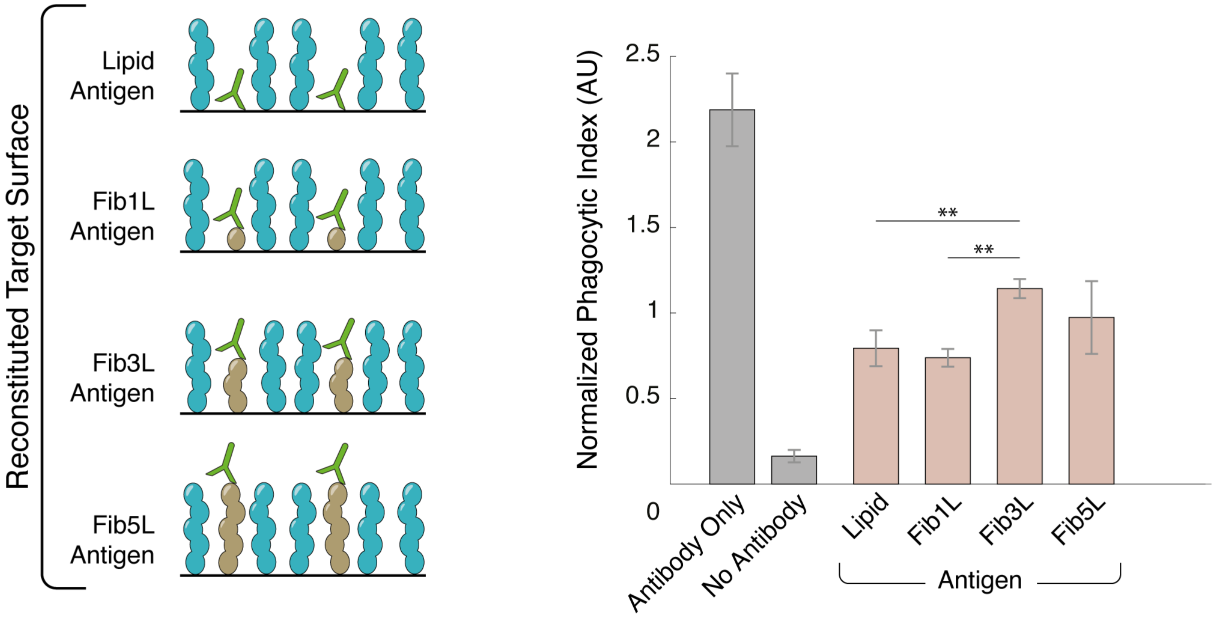


**Fig. S2. Reducing the height difference between bystander proteins and antibody-bound antigens increases phagocytosis.** Phagocytosis was assayed for reconstituted targets displaying different antigen heights in the presence of Fib5L bystander proteins. Antigen densities were fixed at 200 µm$^{-2}$ and bystander proteins were present at the highest possible density (8,000 µm$^{-2}$). When compared to the positive control condition (Antibody Only), phagocytosis is inhibited for all antigen sizes, but maximum phagocytosis occurs for Fib3L antigen ( 10nm) and taller. Error bars represent standard error for >3 replicates of the same condition.

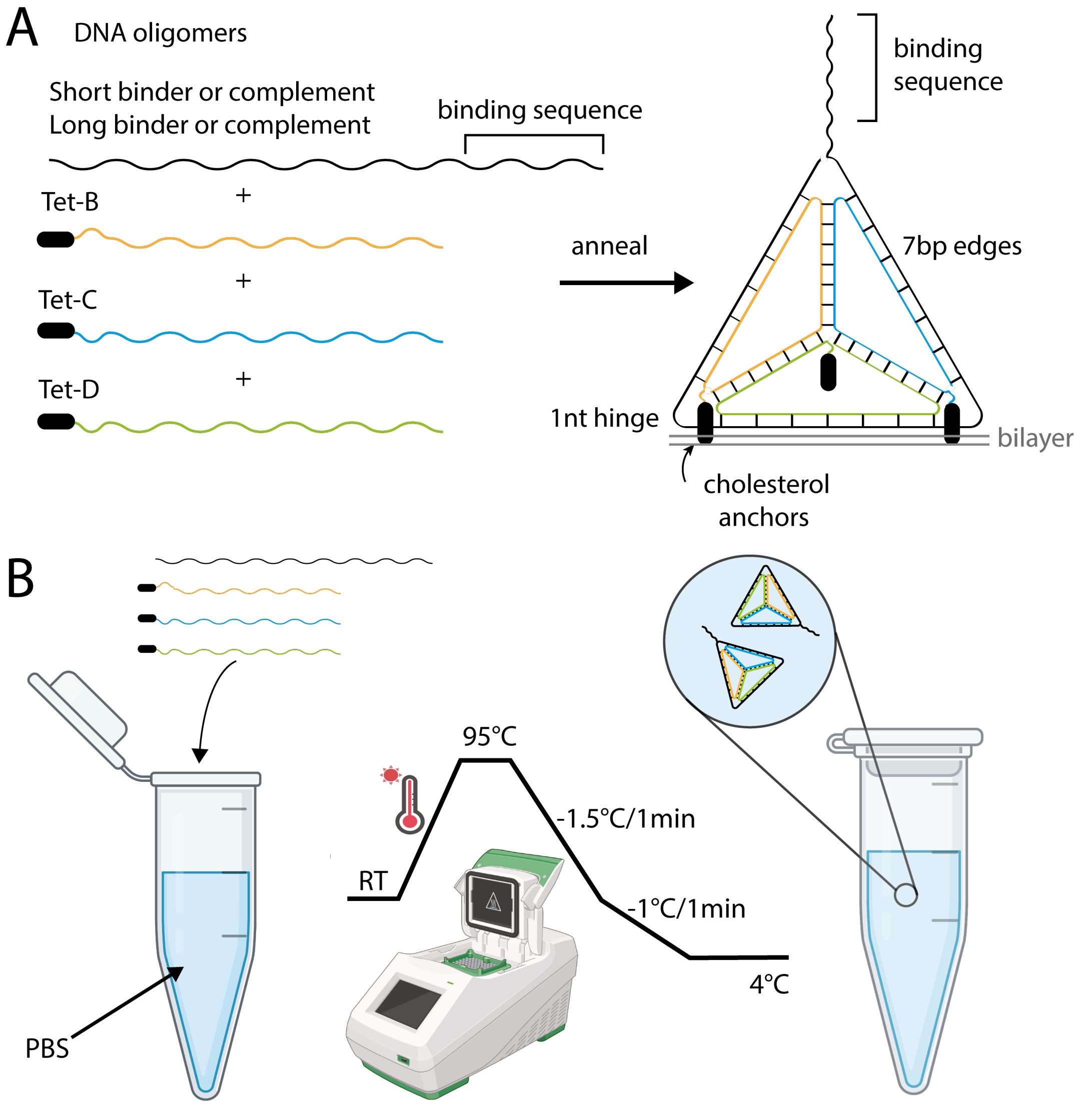


**Fig. S3. Formation and Characterization of DNA Origami-Based Synthetic Binders Used in In Vitro Reconstitution.** **(A)** Tetrahedrons were synthesized by mixing and annealing 4 DNA oligomers. Upon cooling, the strands formed a tetrahedral shape with 7bp edges and one long single-stranded domain protruding from the vertex. **(B)** The individual DNA strands were mixed at equimolar ratios in PBS. They were heated to 95℃ to remove any undesired secondary structure. Gradual cooling was then performed to favor formation of the intended, most thermodynamically stable state—the tetrahedron.

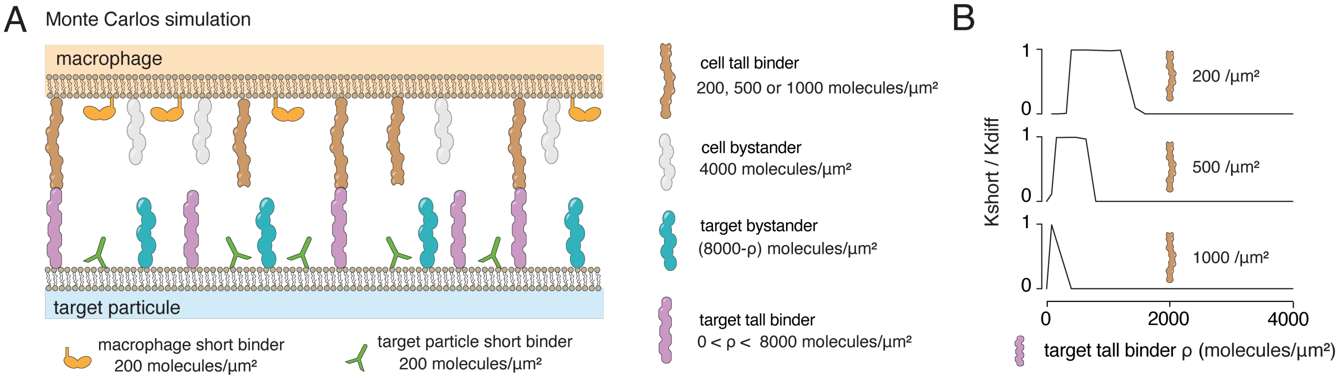


**Fig. S4. Tall binder density modulates short binder kinetics (Monte Carlo simulation) (A)** Monte Carlo simulation of a model macrophage-target interface with bystander molecules and short binders mimicking the FcR-antibody pair. Tall binder molecules, consisting of cell and target components, mimic the PSEL-PSGL1 pair. The densities of short binders (both cell and target) were fixed at 200 molecules/µm$^2$, while cell-side bystanders were maintained at 4000 molecules/µm$^2$. The tall binder density was varied by independently adjusting two parameters: the cell tall binder density (200, 500, or 1000 molecules/µm$^2$) and the density ($\rho$) of target complementary tall binders (0 to 4000 molecules/µm$^2$). To maintain a constant total molecular density of 8000 molecules/µm$^2$ on the target surface, the target bystander density was adjusted accordingly. **(B)** Corresponding kinetic rate of short binder engagement as a function of target tall binder density ($\rho$) demonstrates density-dependent modulation of kinetic enhancement regimes across different cell tall binder densities. For the kinetic enhancement effect to start, a sufficient number of tall binder pairs must form so that the interaction energy ($F_{\text{int}}$) is large enough to lower the activation barrier (see SI text 1B; Steady state kinetics). When cell tall binders are at low density, a high density of target tall binders is needed to reach sufficient probability of forming enough tall binder pairs, which explains why the enhancement effect ($k_{\text{short}}/k_{\text{diff}} \approx 1$) begins at high target tall binder densities (top curve, 200/µm$^2$). Conversely, for high cell tall binder densities (lower curve, 1000/µm$^2$), the critical number of pairs is reached at a lower density of target tall binders. A kinetic enhancement plateau ($k_{\text{short}}/k_{\text{diff}} \approx 1$) persists while the pair density remains sufficient. However, above a critical value, too many tall binder pairs stabilize the intermediate state, maintaining the membranes at a distance (binder pair length) where membrane fluctuations become insufficient for short binder engagement. This causes the engagement kinetics to collapse ($k_{\text{short}}/k_{\text{diff}} \approx 0$). For low cell tall binder density (upper curve, 200/µm$^2$), a high concentration of target tall binders is required to form enough pairs to stabilize the interface at an excessive distance—explaining the wide plateau of kinetic enhancement. In contrast, at high cell tall binder density (lower curve, 1000/µm$^2$), this critical number of pairs is achieved at much lower target tall binder densities, resulting in a narrower kinetic enhancement zone.
